# Spatial proximity determines post-speciation introgression in Solanum

**DOI:** 10.1101/529115

**Authors:** Jennafer A. P. Hamlin, Leonie C. Moyle

**Affiliations:** Department of Plant Biology, University of Georgia, 2502 Miller Plant Sciences, Athens, GA, USA, 30602; Department of Biology, Indiana University, 1001 East Third Street, Bloomington, Indiana, USA, 47405

**Keywords:** Hybridization, Geographic proximity, Mating system variation, Whole-Genome, tomato

## Abstract

An increasing number of phylogenomic studies have documented a clear ‘footprint’ of post-speciation introgression among closely-related species. Nonetheless, systematic genome-wide studies of factors influencing the likelihood of introgression remain rare. Here, we use an *a priori* hypothesis-testing framework, and introgression statistics, to evaluate the prevalence and frequency of introgression. Specifically, with whole genome sequences from 32 lineages of wild tomato species, we assess the effect of three factors on introgression: genetic relatedness, geographical proximity, and mating system differences. Using multiple trios within the ‘ABBA-BABA’ test, we find that one of our factors, geographic proximity, is consistently associated with evidence for recent introgression between species. Of 14 species pairs with ‘proximate’ versus ‘distant’ population comparisons, 12 showed evidence of introgression; in ten of these cases, this was more prevalent between geographically-closer populations. We found no evidence that introgression varies systematically with increasing genetic divergence between lineages or with mating system differences, although we have limited power to address the latter effect. While our analysis indicates that recent post-speciation introgression is frequent in this group, estimated levels of genetic exchange are modest (0.05-1.5% of the genome), so the relative importance of hybridization in shaping the evolutionary trajectories of these species could be limited. Regardless, similar clade-wide analyses of genomic introgression would be valuable for disentangling the major ecological, reproductive, and historical determinants of post-speciation gene flow, and for assessing the relative importance of introgression as a source of evolutionary change.

**IMPACT STATEMENT:** The formation of new species is traditionally viewed as a tree-like branching process, in which species are discrete branches that no longer share an ongoing genealogical connection with other, equally discrete, species. Recently this view has been challenged by numerous studies examining genealogical patterns across entire genomes (all the DNA of an organism); these studies suggest that the exchange of genes between different species (known as ‘introgression’) is much more common than previously appreciated. This unexpected observation raises questions about which conditions are most important in determining whether species continue to exchange genes after they diverge. Factors such as physical proximity, differences in reproductive mechanisms, and time since species shared a common ancestor, might all contribute to determining the prevalence of introgression. But to evaluate the general importance of these factors requires more than individual cases; many species comparisons, that differ systematically in one or more of these conditions, are needed. Here we use whole-genome information from 32 lineages to evaluate patterns of introgression among multiple species in a single, closely related group—the wild tomatoes of south America. We contrast these patterns among pairs of lineages that differ in their geographical proximity, reproductive system, and time since common ancestry, to assess the individual influence of each condition on the prevalence of introgression. We find that only one of our factors—geographical proximity—is consistently associated with greater evidence for recent introgression, indicating that this is largely shaped by the geographical opportunity for hybridization, rather than other plausible biological processes. Our study is one of the first to systematically assess the influence of general ecological and evolutionary conditions on the frequency of post-speciation introgression. It also provides a straightforward, generalizable, hypothesis-testing framework for similar systematic analyses of introgression in groups of other organisms in the future.

## INTRODUCTION

The prevalence of hybridization among species, and the importance of introgression for shaping species evolution, are historically contentious questions (Mallet, 2008, 2005). Although traditionally viewed to be more common among plants (Anderson, 1968; Stebbins, 1970), evidence of hybridization and introgression is emerging for an increasingly broad range of organisms (Mallet et al., 2016). Perhaps the most famous contemporary example involves Neanderthal and modern human lineages (Mallet et al., 2016), in which ~1 – 4% of Neanderthal genome is inferred to have introgressed into some human populations. Quantifying the frequency and amount of introgression is important for understanding the historical dynamics of closely-related lineages, as well as the potential sources of genetic variation that could fuel ongoing evolutionary change. For example, if sufficiently common, gene flow between species could act as a significant source of adaptive loci, as has been observed for mimicry pattern alleles in *Heliconius* butterflies (The Heliconius Genome Consortium et al., 2012). Adaptive introgression is likely to be especially important among recently diverged lineages, where the accumulation of hybrid incompatibilities is not so advanced that it prevents the exchange of unconditionally adaptive loci when lineages come into contact. Nonetheless, the clade-wide prevalence of introgression events, and therefore their relative importance in shaping the evolutionary trajectory of close relatives, is only now beginning to be assessed (Folk et al., 2018).

From a genomic perspective, introgression leaves a detectable ‘footprint’: introgressed regions show distinctive patterns of historical relatedness that differ from non-introgressed regions, because they are most closely related to the donor species rather than the recipient genome in which they are found (Payseur and Rieseberg, 2016). Accordingly, genome-wide data is ideal for characterizing the prevalence of hybridization because it provides a detailed picture of phylogenetic relationships at loci across the genome, including in genomic regions that show patterns of relatedness inconsistent with the species as a whole. Beyond the human and butterfly examples, genome-wide data has been used to infer past introgression events among species in groups as diverse as *Saccharomyces* yeast (Morales and Dujon, 2012) *Anopheles* mosquitoes (Fontaine et al., 2015), wild tomatoes (Pease et al., 2016), and *Drosophila* (Turissini and Matute, 2017). However, while revealing the extent and timing of gene flow events is interesting in individual cases, there are few tests of the generality of introgression across whole groups of closely related species, including whether it systematically varies in frequency or extent under different biological conditions.

Some of the factors that could influence the frequency of hybridization and subsequent introgression include phylogenetic relatedness (i.e., genetic distance), geographical proximity, and biological factors that affect the likelihood and direction of reproductive events, such as differences in mating system. In the first case, because the strength of reproductive isolation is expected to accumulate with the amount of time since lineages diverged (Coyne and Orr, 1989), more genetic exchange might be expected to occur between more closely-related species, with diminishing rates accompanying increasing lineage differentiation. Second, genetic exchange is more likely to occur among species in close geographic proximity, where they can potentially come into physical and therefore reproductive contact (Harrison, 2012). Determining the level of spatial proximity that allows gene exchange can be challenging, as it likely depends upon numerous biological factors (e.g., dispersal mechanisms) and abiotic factors (e.g., physical barriers to dispersal). Nonetheless, a general expectation is that hybridization is more likely when biological or abiotic factors do not hinder contact between lineages. For example, numerous hybrid zone studies demonstrate that the proportion of individuals with mixed ancestry usually decreases with geographic distance from the hybrid zone (Harrison and Larson, 2014, 2016).

Third, factors that specifically influence the timing and success of reproductive events are also expected to influence the likelihood of hybridization and introgression. For example, mating system variation can influence introgression, either directly by affecting the likelihood of successful mating between species or indirectly by influencing the longer-term likelihood that introgressed loci will persist in the recipient lineage. In the first instance, mating system differences can cause predictable asymmetries in the success of initial crosses between species. This can occur either via differences in the size or shape of reproductive organs that can lead to asymmetric mechanical isolation between lineages (e.g. where outcrossing species can fertilize inbreeding species, but not vice versa; Brothers and Delph, 2017; Levin, 1978) or—especially in plants—via differences in the presence/absence of genetically determined self-incompatibility systems, whereby pollen from self-incompatible species can fertilize ovules of self-compatible species, but self-incompatible plants actively reject pollen from self-compatible species (e.g. in *Nicotiana* (Anderson and de Winton, 1931), *Petunia* (Mather and Edwardes, 1943) and *Solanum* (McGuire and Rick, 1954)). In both mechanical and active-rejection cases, outcrossing species are more likely to donate alleles to more inbreeding species compared to the reciprocal direction of gene flow, reducing the potential for gene flow specifically between species with unalike mating systems. Similarly, the longer-term likelihood that introgressed loci will persist in recipient lineages can vary based on the mating system of the donor and recipient lineages, because mutational load and the efficacy of selection is expected to differ between species with histories of more or less inbreeding and different effective population sizes (*N_e_*) (Busch, 2005; Charlesworth et al., 1990; Harris and Nielsen, 2016; Juric et al., 2016; Lande and Schemske, 1985). In particular, introgression from outbreeding to inbreeding populations should be especially disfavored both because donor alleles are expected to have stronger deleterious fitness effects (due to genetic load that can persist in outbreeders) and because the smaller *N_e_* recipient population is less effective at disassociating these from other non-deleterious loci before they are purged (Brandvain et al., 2014; Ruhsam et al., 2011). In comparison, the exchange of alleles between lineages with similar mating systems should be less constrained by these considerations. In general, then, no matter whether affected by initial crossing differences (from mechanical or active rejection asymmetries) or differences in the historical factors determining genetic load and effective population size, gene flow between lineages that differ in their mating system might be expected to be more constrained than gene flow between lineages with similar mating systems.

While these factors are expected to influence the rate and likelihood of gene flow between species, there are few systematic tests of their general importance in shaping the prevalence of post-speciation introgression. Here we use whole genome data to systematically evaluate the effects of genetic distance, geographical proximity, and mating system differences, on genome-wide patterns of post-speciation introgression across a closely related clade of species. To assess introgression, we use the ‘ABBA-BABA’ test (also known as the D-test; Durand et al., 2011; Green et al., 2010). This test detects introgression by comparing the frequency of alternate ancestral (“A”) and derived (“B”) allele patterns among four taxa, where the species tree has the allele pattern BBAA (Figure 1). In the absence of gene flow, the alternate minority patterns of ABBA and BABA should be approximately equally frequent, as they have an equal chance of either coalescence pattern under incomplete lineage sorting (ILS; Durand et al., 2011). In comparison, an excess of ABBA patterns indicates gene flow between lineage P2 and P3, and excess BABA indicates gene flow between lineage P1 and P3 (Figure 1).

**Figure 1.**
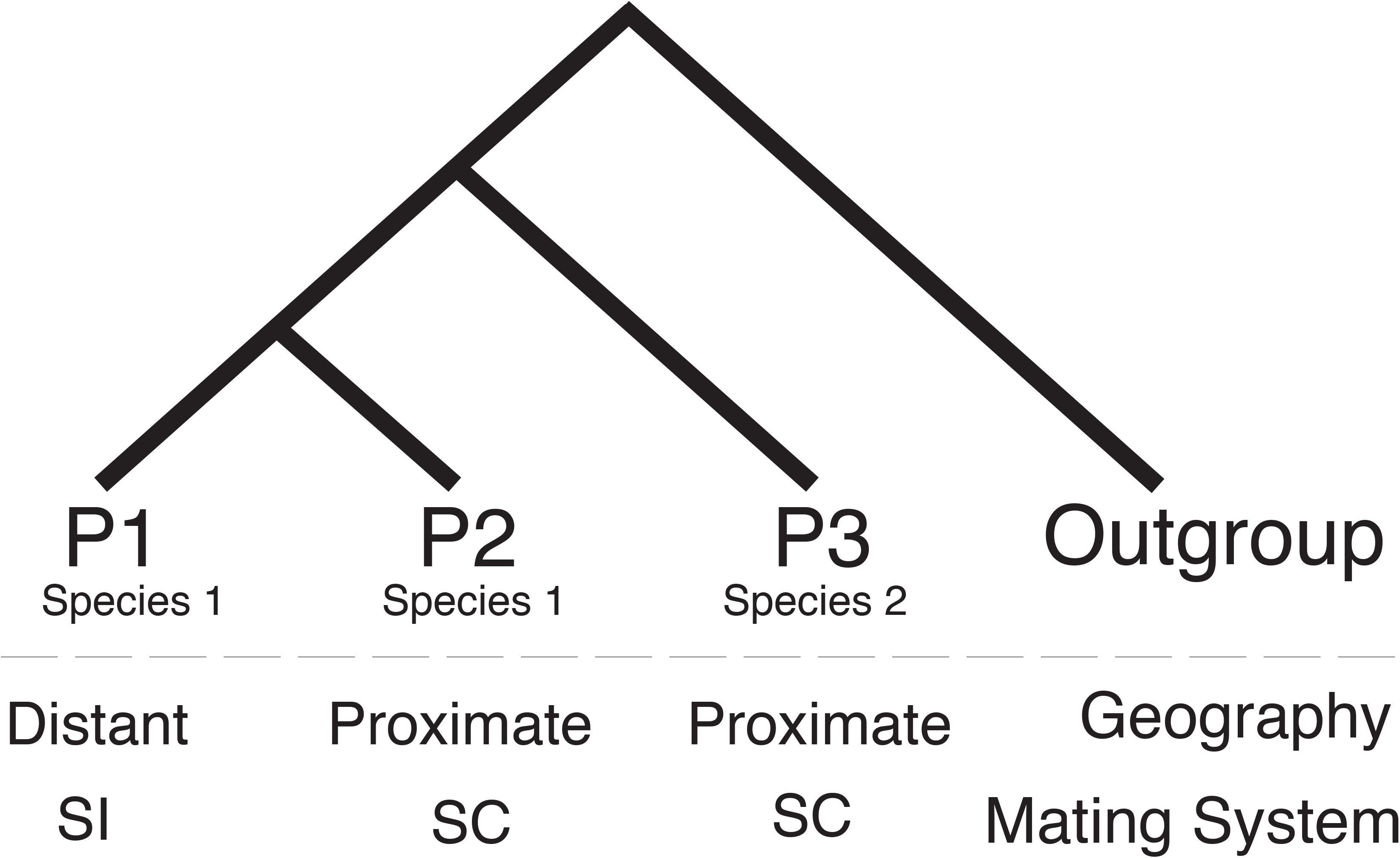
Structured ABBA-BABA tests to evaluate *a priori* hypotheses about the expected prevalence of introgression due to geographical proximity and/or lineage differences in mating system. For example, if introgression occurs more frequently between geographically closer accessions, more minority topologies should support a closer relationship between P2 and P3, compared to P1 and P3, and the genome-wide mean D-statistic is expected to be positive.

Importantly, the structure of the ABBA-BABA test allows us to test *a priori* hypotheses about the expected prevalence of introgression in multiple parallel comparisons. In particular, we can intentionally construct tests of a specific introgression hypothesis by consistently positioning taxa in the P1, P2, and P3 positions in a 4-taxon tree, so that P2 and P3 are always predicted to experience more introgression than P1 and P3 (Figure 1). For example, if geographic proximity *per se* affects the amount of introgression between two species, in a case where P1 and P2 are populations of one species but P2 is more geographically proximate to populations of a second species (P3), then our *a priori* expectation is that elevated introgression will be observed as an excess of ABBA (i.e. evidence of gene flow between P2 and P3) rather than BABA (gene flow between P1 and P3; Figure 1). Multiple different four-taxon tests with the same structure can then be used to evaluate whether geographically proximity is systematically more frequently associated with evidence of post-speciation introgression. A similar structure can be used to test the *a priori* expectation that gene flow is expected to be more frequent between lineages with shared versus different mating systems. More generally, 4-taxon tests that involve increasing evolutionary divergence between the P1/P2 and P3 lineages, can be used to evaluate evidence that introgression is on average more prevalent between more closely related taxa.

Here we use this *a priori* hypothesis-testing framework to assess the prevalence and frequency of introgression among wild tomato lineages (*Solanum* section *Lycopersicum*) depending upon a) geographical proximity, b) differences in mating system, and c) evolutionary distance. In addition to high-quality, whole-genome data for multiple wild genotypes of known provenance (Aflitos et al., 2014; Lin et al., 2014), this group has well-curated historical records of geographic coordinates for hundreds of wild populations from across the clade. Moreover, prior evidence of introgression events between specific lineages (e.g. Beddows et al., 2017; Pease et al., 2016), and the ability to generate F1 and later-generation hybrids in the greenhouse (e.g., Moyle, 2008; Rick, 1979), indicate the possibility that introgression could shape genomes in this group. Using whole-genome data from 32 closely related accessions across 12 species of wild tomato, our goal here was to systematically test hypotheses about the prevalence of introgression, to make general inferences about the role and importance of particular factors in the frequency of cross-species hybridization, and to begin to assess the potential importance of introgression in shaping genome content and evolutionary trajectories in this clade.

## METHODS

### Sequencing data and mapping to reference genome

Our analyses used data from three whole genome-sequencing projects (Aflitos et al., 2014; Hardigan et al., 2016; Lin et al., 2014). Specifically, we obtained raw reads either as fastq or SRA files for genomes of 32 wild *Solanum* individuals from the tomato clade (*Solanum* section *Lycopersicum*), each from a different accession (historical population collection), along with *Solanum tuberosum* (potato; Hardigan et al., 2016) which we used as the outgroup in all comparisons (Supplemental Table 1). To combine data from the different sequencing projects, we trimmed and re-mapped raw reads back to the reference genome of domesticated tomato, *S. lycopersicum* version 2.50 (The Tomato Genome Consortium, 2012), using standard practices for mapping and quality-filtering (see Supplemental Text).

**Table 1.** Introgression statistics for each analyzed trio (4-taxon test), based on data from 100kb windows with > 20 informative sites. Trios in bold have D statistics that are significantly different than zero after Bonferroni correction. The order in which each species accession is listed corresponds to (P1, P2, P3). In all instances, we use the potato genome (*S. tuberosum*) as the outgroup.

| Group | Accessions | MeanD | StnDev | SE | C95_lwr | C95_upr | Chi-square | p.value |
| --- | --- | --- | --- | --- | --- | --- | --- | --- |
| *a. Geographic Trios\** |  |  |  |  |  |  |  |  |
| gal.gal.che | LA1044.LA0483.LA0746 | -0.0277 | 0.4947 | 0.0430 | -0.1120 | 0.0566 | 0.97 | 1 |
| **arc.arc.pim** | **LA2172.LA2157.LA2147** | **0.0328** | **0.5374** | **0.0072** | **0.0186** | **0.0470** | **1666.13** | **2.2e-16** |
| **pim.pim.neo** | **LA1375.LA1246.LA2133** | **-0.0892** | **0.5902** | **0.0073** | **-0.1036** | **-0.0749** | **102.83** | **5.51E-23** |
| pim.pim.chi | LA1582.LA1933.LA1969 | -0.0100 | 0.5168 | 0.0094 | -0.0285 | 0.0085 | 2.82 | 1 |
| **pim.pim.cor1** | **LA0400.LA1269.LA0118** | **0.0162** | **0.5075** | **0.0080** | **0.0005** | **0.0318** | **10.30** | **1.82E-02** |
| **pim.pim.cor2** | **LA1617.LA1521.LA0118** | **0.0998** | **0.6441** | **0.0082** | **0.0838** | **0.1158** | **126.44** | **3.44E-28** |
| **pim.pim.per1** | **LA1595.LA1341.LA1278** | **0.0047** | **0.5091** | **0.0070** | **-0.0090** | **0.0184** | **21.44** | **5.10E-05** |
| **pim.pim.per2** | **LA1617.LA1269.LA1278** | **0.0828** | **0.6145** | **0.0081** | **0.0669** | **0.0987** | **85.52** | **3.21E-19** |
| **pim.pim.hab** | **LA0417.LA0442.LA1777** | **0.0421** | **0.5683** | **0.0072** | **0.0280** | **0.0561** | **96.14** | **1.49E-21** |
| **pim.pim.pen** | **LA1245.LA1269.LA1272** | **0.0683** | **0.5357** | **0.0091** | **0.0504** | **0.0862** | **72.93** | **1.87E-16** |
| **arc.arc.hab** | **LA2172.LA2157.LA1718** | **-0.0154** | **0.4198** | **0.0074** | **-0.0299** | **-0.0009** | **22.35** | **3.17E-05** |
| **hab.hab.neo** | **LA1777.LA1718.LA2133** | **0.0483** | **0.5144** | **0.0072** | **0.0342** | **0.0624** | **47.32** | **8.42E-11** |
| **hab.hab.cor** | **LA0407.LA1777.LA0118** | **0.0579** | **0.4796** | **0.0072** | **0.0439** | **0.0720** | **161.13** | **8.95E-36** |
| **hua.hua.hab** | **LA1983.LA1365.LA1718** | **0.0532** | **0.4348** | **0.0076** | **0.0383** | **0.0680** | **364.90** | **3.37E-80** |
| *b. Mating System Trios* |  |  |  |  |  |  |  |  |
| **arcSl.arcSC.pimSC** | **LA2172.LA2157.LA0373** | **0.04356** | **0.531** | **0.282** | **0.007** | **0.030** | **1999.06** | **2.2e-16** |
| **arcSl.arcSC.pimSC2** | **LA2172.LA2157.LA0400** | **0.0264** | **0.547** | **0.008** | **0.011** | **0.042** | **736.04** | **1.31E-161** |
| **habSl.habSC.pimSC** | **LA1777.LA0407.LA0373** | **-0.03551** | **0.508** | **0.258** | **0.006** | **-0.048** | **42449.7** | **2.2e-16** |
| **habSl.habSC.pimSC2** | **LA1777.LA0407.LA0400** | **-0.0357** | **0.560** | **0.007** | **-0.050** | **-0.021** | **5526.61** | **2.2e-16** |
| **perSl.perSC.pimSC** | **LA1278.PI128650.LA0373** | **0.14860** | **0.554** | **0.307** | **0.008** | **0.134** | **5456.4** | **2.2e-16** |
| **perSl.perSC.pimSC2** | **LA1278.PI128650.LA0400** | **0.1526** | **0.562** | **0.010** | **0.133** | **0.173** | **8.79** | **9.10E-03** |

StnDev = standard deviation; SE = standard error of mean; C95\_lwr and C95\_upr = Lower and upper 95% confidence intervals, respectively, Chi-square = Chi-squared goodness-of-fit test value. \* For geographic trios with non-overlapping accessions, see supplemental table 7.

### Hypothesis testing with the D-statistic

We used the ABBA-BABA test to assess evidence for the presence and directionality of gene flow in a set of 4-taxon tests. The results of each ABBA-BABA test can be expressed in terms of Patterson’s *D*-statistic, calculated as (#ABBA - #BABA) / (#ABBA + #BABA) for all biallelic sites in the multiple sequence alignment (Durand et al., 2011; Green et al., 2010). The D-statistic therefore summarizes both the magnitude of introgression and the specific pair of taxa that are exchanging alleles; positive values of D indicate P2 and P3 are exchanging more alleles (an excess of ABBA) and negative values indicate more gene exchange between P1 and P3 (an excess of BABA). We used multiple replicate 4-taxon tests to evaluate three *a priori* expectations:

1. Post-speciation introgression is more prevalent between geographically closer versus more distant lineages. Four-taxon tests were structured so that P1 and P2 were taken from populations of the same species, but P2 was spatially closer to the P3 species and P1 was more distant (Figure 1). In this case, we expect a systematic excess of positive values of the resulting *D*-statistics. Species comparisons and specific P1, P2, and P3 accessions were identified for these tests based on known species ranges and geographical locations of the sequenced accessions (see Supplemental text for our specific criteria). Because these analyses are constrained by the available sequenced genotypes, the actual geographic distances involved vary broadly between 4-taxon tests (Supplemental Table 2), so that this analysis is an imperfect reflection of close spatial proximity; however, the structure of each test means we are still able to systematically compare the effect of greater (‘proximate’) versus less (‘distant’) geographic proximity on detected patterns of introgression.
2. Post-speciation introgression is more prevalent between lineages that share mating system. Here, four-taxon tests were structured so that P1 and P2 were again taken from populations of the same species, but P2 was self-compatible (SC) and P1 was self-incompatible (SI); in every test, P3 was the same accession of an SC species. In this case, we also expect a systematic excess of positive *D*-statistics. Within our dataset there are only three species for which we had whole genome sequence data from both SI and SC accessions: *S. arcanum*, *S. habrochaites*, and *S. peruvianum*.
3. Post-speciation introgression is more prevalent between lineages that are more closely related. In this case, we expect that the estimated magnitude of D should decrease as evolutionary (genetic) distance between (P1, P2) and P3 increases within the 4-taxon test. We calculated pairwise genetic distance for each comparison by taking the average genetic distance for the two comparisons within the focal trio (i.e. P1 with P3 and P2 with P3), based on genome-wide site differences between accessions. In this case, there is no expectation of either positive or negative *D*-values, as P1 and P2 are equally closely related to P3. Instead, the comparison is among *D*-values taken from different 4-taxon combinations.

The supplemental text provides a more detailed description of how these factors were defined and determined for individual 4-taxon tests.

### Calculating and analyzing D-statistics

Using the program *mvftools* (Pease and Rosenzweig, 2018), we estimated D-statistics for consecutive 100kb windows across each of the 12 unique chromosomes of *Solanum*, and then calculated the genome-wide average of D across all of these windows. 100 kb windows were defined based on the domesticated tomato genome, resulting in 8,015,000 total windows (0.8015 gigabases or ~ 80% of the genome). For all analyses reported in the main text, we used only 100 kb windows that had >20 variable sites (~4200 windows on average per test, depending upon the specific trio analyzed; Supplemental Table 3); this eliminated windows in which no or few single nucleotide polymorphisms (SNPs) were found, as these have low power to accurately estimate *D*. For completeness, we also performed analyses on *D* estimated from all 100 kb windows in the genome and found the same qualitative results as the higher power dataset (Supplemental Table 4). For each four-taxon test, we performed a chi-squared goodness-of-fit test, to determine if our genome-wide estimate of mean *D* was significantly different than zero. Across all four-taxon tests, we determined whether the number of tests supporting a higher incidence of introgression in the predicted direction was greater than expected by chance, using a sign test. All statistical analyses were performed in RStudio version 1.1453 (2015).

To determine if post-speciation introgression is associated with quantitative differences in geographical proximity between populations of two species, for every trio we also determined the geographic distance (in km) between P1 and its nearest P3 accession, and between P2 and its nearest P3 accession (using our georeferenced location data for all population accessions, see supplemental text), to generate an estimate of their relative proximity to any population from P3 (i.e. the difference between these two distances; Supplemental Table 5). For all four-taxon tests, we regressed our mean genome-wide estimate of *D* on this relative geographic distance estimate (Supplemental Figure 1). To determine if evidence for post-speciation introgression is more prevalent for recently diverged lineages, we evaluated the association between the absolute values of our estimates of genome-wide *D* and genome-wide genetic distance, using regression across all four-taxon tests.

Finally, to provide a general estimate of the fraction of the genome inferred to have come from introgression in each trio, we calculated the number of varying sites that support evidence of differential introgression (i.e. the excess of sites that support one minority topology over the other, or (|ABBA sites – BABA sites|) and expressed these as a proportion of total variable sites in this trio (i.e. |ABBA – BABA| / (ABBA + BABA + BBAA)). These estimates were calculated for all four-taxon combinations included in our analyses, and used only data from 100 kb windows with >20 SNPs (Supplemental Table 6). While these estimates are not precise measures of the proportion of the genome that has experienced introgression (for example, they could underestimate total introgression if there has been gene flow both between P1 and P3, and between P2 and P3), they provide a set of rough global estimates with which to compare different four-taxon tests. These estimates were regressed onto mean genetic distance for all trios to evaluate whether the amount of inferred introgression between species was associated with the time since their divergence (Supplemental Figure 2).

**Figure 2.**
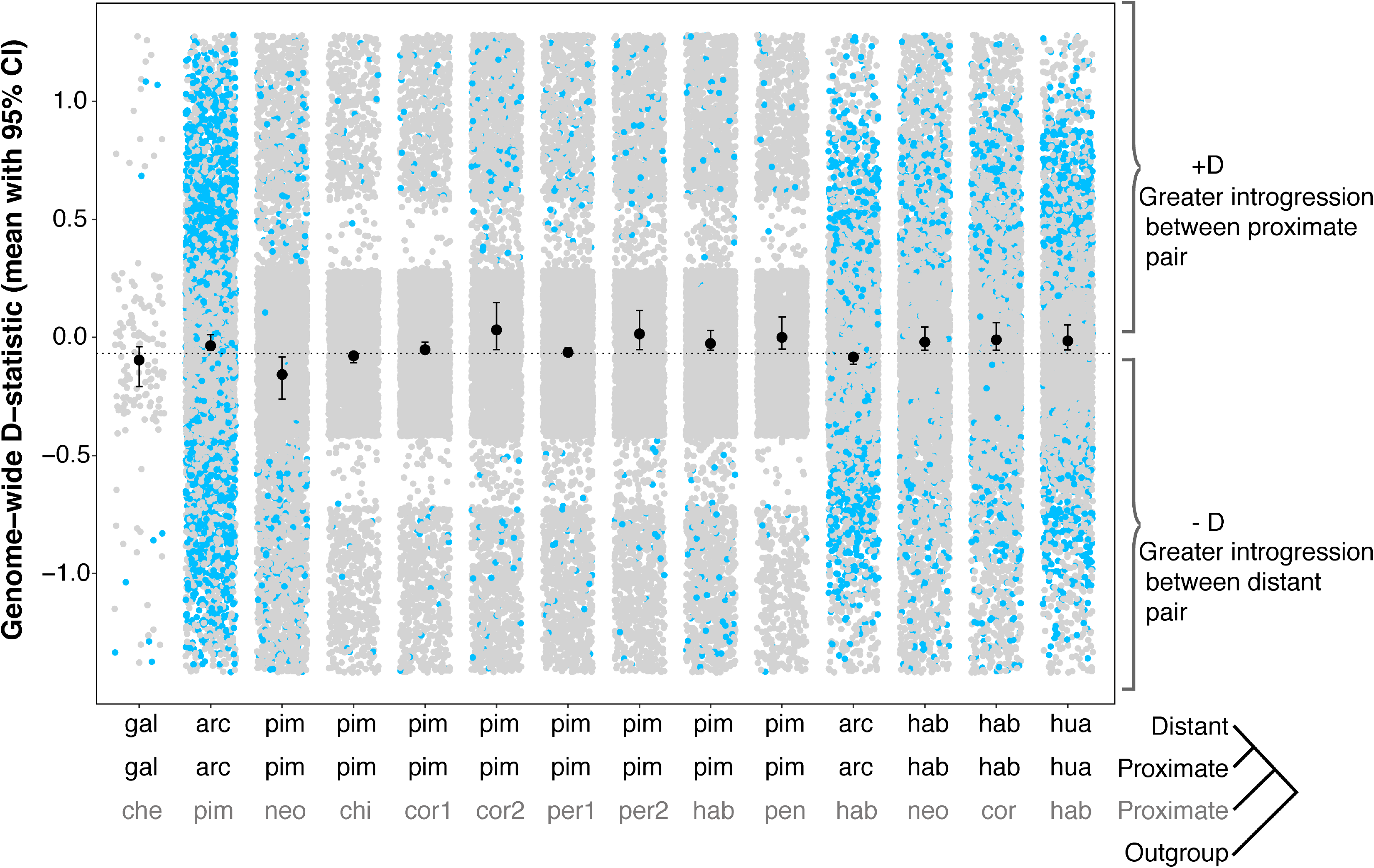
Effect of geographical proximity on introgression. For each geographic trio, the plot shows genome-wide mean *D* values and 95% confidence intervals, as well as *D*-statistic estimates from individual 100kb windows (gray circles: individual window *D* values not significantly different than zero; blue circles: individual window *D* values significantly different than zero). Accessions in each trio are listed in Table 1. For all trios, our outgroup is potato (*S. tuberosum*).

## RESULTS

We generated genome-wide D estimates for 17 four-taxon combinations, 14 that could address the effects of geographical proximity on introgression, and three addressing the effects of mating system variation. All 17 combinations were used to assess the effect of genetic distance. Across all 17 tests, the genome-wide average estimates of *D* ranged from −0.08 to 0.15, based on 100kb windows with >20 variable sites (Table 1; estimates of *D* from all windows were similar, Supplemental Table 4). The fraction of the genome estimated to be differentially introgressed ranged from 0.05 – 1.5 (Supplemental Table 6).

For our tests of geographical proximity, of 14 testable four-taxon combinations, ten had average *D* values significantly greater than zero—indicating that our geographically closer lineages (P2 and P3) share a higher proportion of sites—whereas two were significantly less than zero (chi-squared goodness-of-fit tests, with Bonferroni correction; Figure 2, Table 1a). A two-sided sign-test indicated that the number of significantly positive versus negative mean values of *D* was different (p = 0.038), consistent with a systematic excess of introgression between species pairs when their populations are geographically closer versus more distant. Because there is non-independence in our dataset (that is, some individual accessions/genome sequences are used in more than one four-taxon test), we evaluated the influence of this non-independence by paring our dataset down to trios (of P1, P2, P3) that only used unique accessions. Of our twelve trios with significant positive or negative *D* statistics, we could evaluate combinations of up to six different trios that shared no accessions in common. In all eight unique alternative combinations of six trios, five tests had a positive D and one had a negative D; that is, we found evidence for an excess of introgression consistently more frequently when populations of a species pair were geographically closer versus more distant. This directionality is non-significant in each of the reduced datasets (p = 0. 2188; Supplemental Table 7) as the two sided sign test is underpowered to detect a systematic difference in direction when n=6. Finally, across all 14 four-taxon tests, mean genome-wide *D* was not significantly associated with the relative geographical proximity of the P1 versus P2 population to the closest population from the P3 species (R-squared = 0.055, p-value = 0.42, Supplemental Figure 1a), suggesting that, while geographical proximity influences the possibility of introgression, other factors likely determine the quantitative amount of introgression that occurs.

For our evaluation of mating system effects, of three testable trios, two genome-wide mean *D* values were significantly positive and one was significantly negative, regardless of the specific accession used as the P3 taxon in our tests (Table 1; Figure 3). Note that relative geographical proximity does not explain the sign of *D* in any of these cases; for the two instances where *D* was positive (P1 and P2 from *S. arcanum* or from *S. habrochaites*, respectively: Supplemental Table 5), our SI accessions are geographically closer to a P3 accession compared to their conspecific SC accessions, and the opposite is observed in the case where mean *D* was negative. In addition, mean genome-wide D is not negatively associated with the relative difference in geographic proximity for these mating system trios (Supplemental Figure 1b). Overall, then, our evidence in support of mating system effects is equivocal: 2 of 3 tests support the expectation that introgression is more prevalent between species pairs when their populations share the same mating system, but one does not.

**Figure 3.**
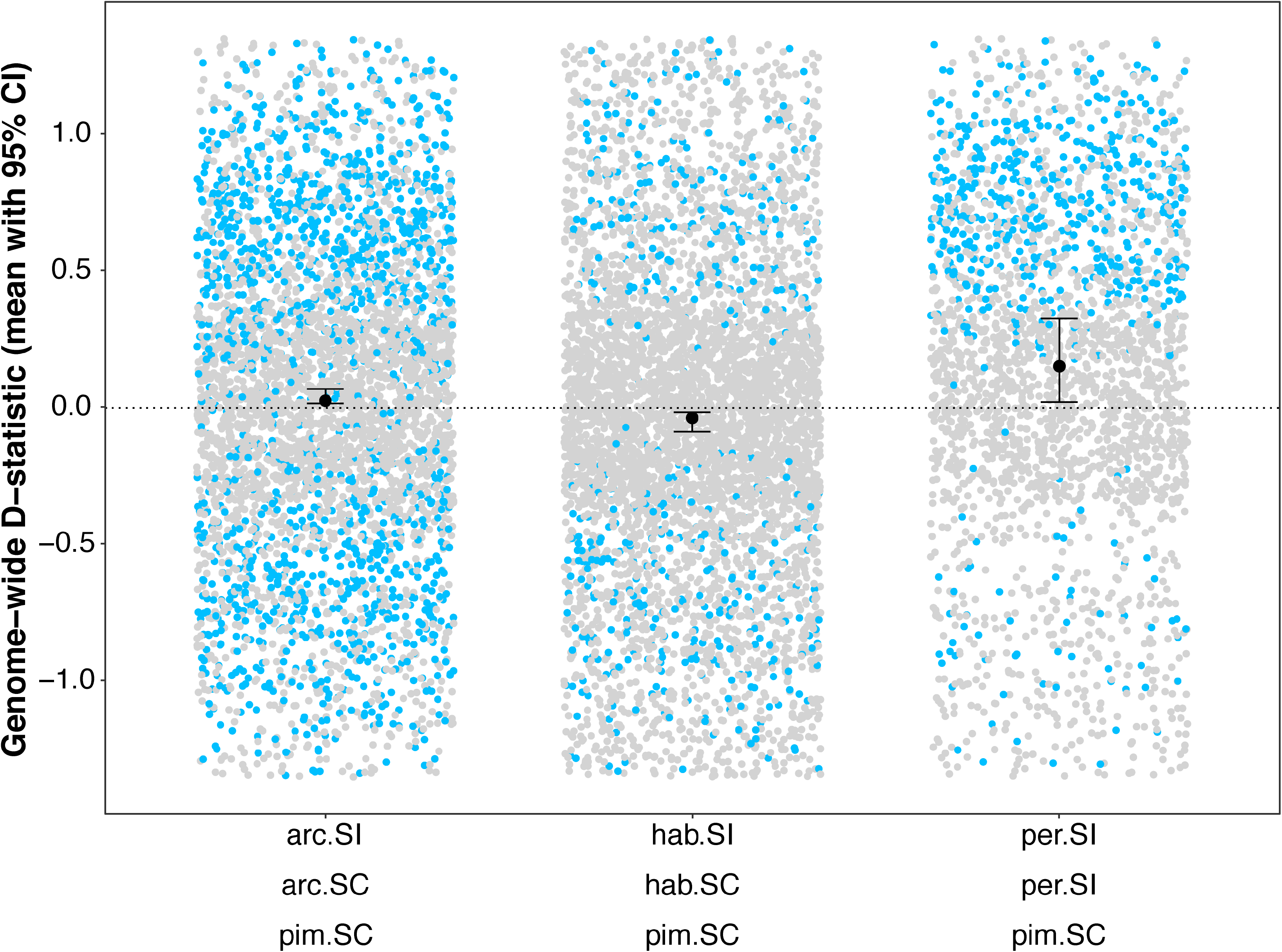
Effect of mating system differences on the observed direction of introgression. For each mating system trio, the plot shows genome-wide mean D and 95% confidence intervals, as well as *D*-statistic estimates from individual 100kb windows (gray circles: individual window *D* values not significantly different than zero; blue circles: individual window *D* values significantly different than zero). Here the P3 position is occupied by the same accession of *S. pimpinefollium* (LA0373) for all three comparisons.

Finally, we detected no association between quantitative values of D and increasing evolutionary divergence between P1/P2 and P3 species (for 14 geographic trios: R-squared = 0.021, p-value = 0.62; for all 17 trios: R-squared = 0.012, p-value = 0.68, Figure 4 and Supplemental Table 8), suggesting little evidence that the propensity for introgression is determined by variation in evolutionary divergence among the species pairs examined here. Similarly, the estimated amount of differential introgression (as a proportion of all variable sites; 0.05%-1.5%) is unrelated to the mean genetic distance between the focal species in each trio (R-squared = 0.0004, p-value = 0.938, Supplemental Figure 2).

**Figure 4.**
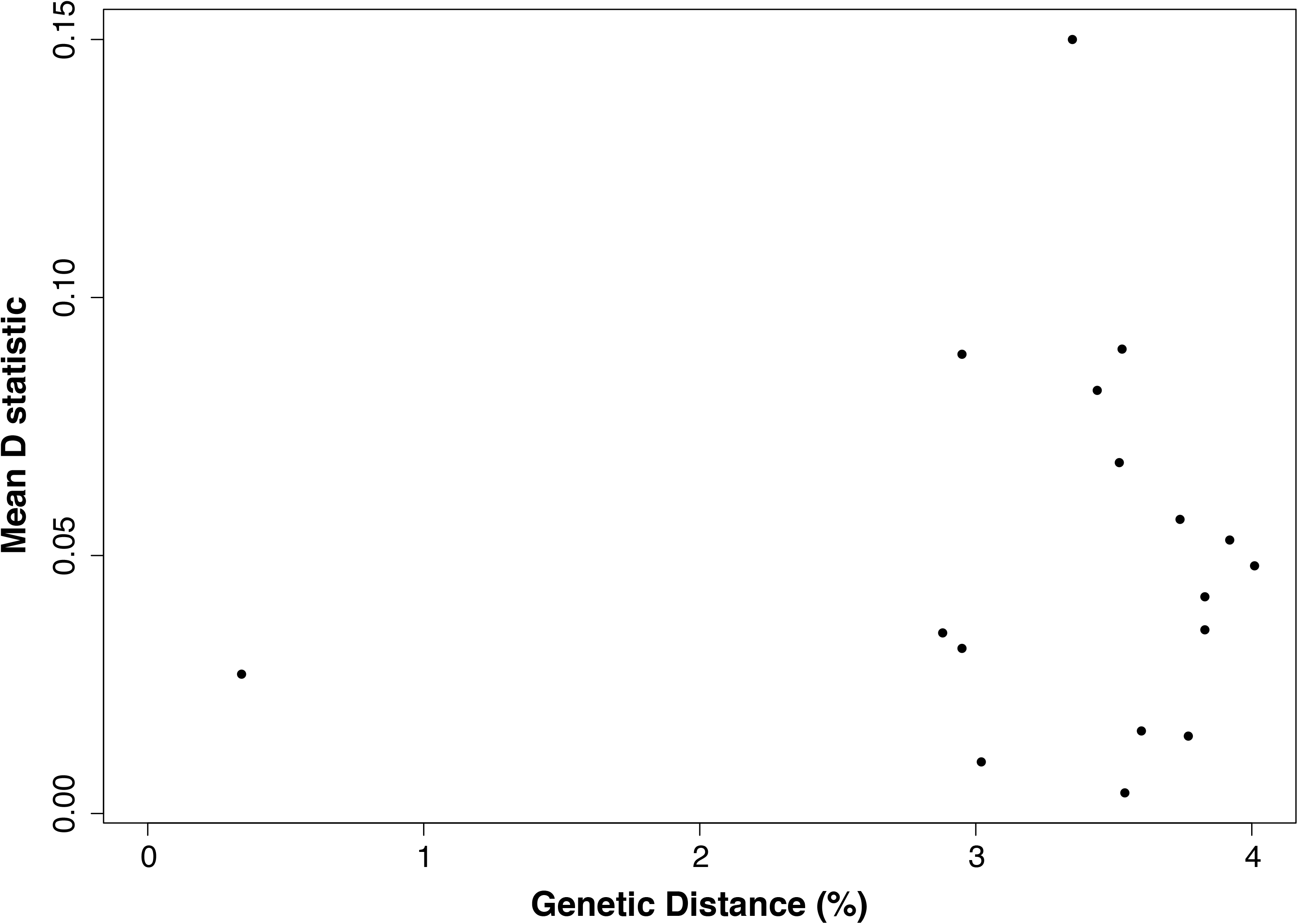
The relationship between genome-wide mean *D* and the average genetic distance (% divergence across all sites) between P1/P2 and P3 species for 14 trios used in geographic tests (R-squared = 0.012, p-value = 0.68).

## DISCUSSION

The prevalence of introgression is one pattern emerging from contemporary genome-wide studies in many groups of closely related species, including in groups not traditionally associated with post-speciation gene flow. However, there have been few attempts to systematically assess the influence of different factors in shaping the frequency and extent of this gene flow. Here we used directionally structured, four-taxon ABBA-BABA tests to examine the influence of three factors—genetic distance, geographical proximity, and mating system differences—on genome-wide patterns of introgression among wild tomato species. We found that recent introgression was commonly detected among these species, that consistent patterns of post-speciation introgression depend largely on geographical proximity rather than the other two factors, and that the estimated fraction of the genome differentially introgressed between species was modest. These findings have interesting implications for interpreting the contexts in which introgression might play the greatest role in shaping evolutionary trajectories in this and other similar clades, and for assessing the potential contribution of introgression to adaptive phenotypic evolution.

### Recent introgression occurs frequently but is modest in scope among wild tomatoes

Our analysis indicates that, among wild tomato lineages, post-speciation gene exchange is prevalent: of 17 total four-taxon tests across all our analyses, 15 had mean *D* values significantly different from zero. Prior studies have detected evidence for introgression among specific wild tomato lineages (Beddows et al., 2017; Pease et al., 2016), and our findings expand and illuminate these observations in several key respects. First, our analyses preferentially assessed evidence for recent, rather than more ancient (Pease et al., 2016), introgression events because in every case we contrasted populations (P1 and P2) from a single species when looking for evidence of introgression with a second species (P3). Accordingly, any inferred introgression must have occurred after the evolutionary split of these two (P1 and P2) conspecific populations. Despite this, we find repeated evidence that populations from different species have exchanged genes recently, including species that are estimated to have diverged >2 million years ago (e.g., *S. pimpinellifolium* and *S. pennellii*; Pease et al., 2016).

Our results suggest there is broad potential for cross-species hybridization across the clade, a finding consistent with other observations that indicate premating isolation is likely to be incomplete between lineages in nature: all species share general floral morphology (rotate, yellow, five-petaled flowers), all are buzz-pollinated, and multiple species pairs are found in sympatry (Rick, 1950). Nonetheless this finding is intriguing as few natural hybrids have been observed in the wild in this group (Taylor, 1986), and some of these species are known via crossing and genetic studies to express moderate to strong postmating and postzygotic reproductive isolation under lab conditions (Hamlin et al., 2017; Moyle and Nakazato, 2010, 2008). These later-acting barriers might be important in limiting the amount of introgression that results from hybridization events. Indeed, a second general observation of our analysis is that despite evidence for relatively frequent hybridization, the amount of the genome exchanged between species is likely to be limited: the proportion of the genome estimated to be differentially exchanged between species is on the order of 0.05-1.5% of all variable sites (Supplementary Table 8). Further, when D-statistics are examined chromosome-by-chromosome within each four-taxon test (Supplementary Figure 3), in most cases introgression is inferred on some chromosomes but not others; this variation among chromosomes might be due to variation in the presence of loci contributing to reproductive isolation. Overall, the amount and distribution of inferred introgression suggests that current species reproductive barriers are sufficiently incomplete to allow detectable recent introgression among diverged species in the field, but also that genomes are not completely or uniformly porous to gene flow among lineages, even in cases where there is an opportunity for gene exchange.

### Introgression frequency varies with spatial proximity between species pairs, rather than overall genetic relatedness or mating system differences

Importantly, the structure of our analyses also allowed us to explicitly evaluate the influence of several factors on these detected patterns of introgression. We found that repeated patterns of recent post-speciation hybridization were consistently associated with only one of our factors: geographical proximity. This suggests that the propensity for post-speciation gene exchange is most often dependent on the simple opportunity for reproductive contact—geographic proximity *per se* between populations of different species allows greater gene flow. In contrast, although there were few comparisons available, we found that mating system differences were not consistently associated with reduced (or elevated) patterns of introgression; two tests were consistent with our *a priori* prediction (including the strongest case) but one test indicated more introgression between lineages that differed in mating system. While we currently have too few four-taxon tests to draw definitive conclusions, these idiosyncratic observations could be due to additional biological factors. For example, the transition to self-compatibility within each of our polymorphic (SI/SC) species might be too recent to yet see consistent differences in crossing asymmetry or genetic load that are expected between long-standing SC versus SI populations. Regardless, because there are clear predictions about associations between mating system, genetic load, the efficacy of selection, and the propensity and direction of introgression (Busch, 2005; Charlesworth et al., 1990; Harris and Nielsen, 2016; Juric et al., 2016; Lande and Schemske, 1985)—some of which appear to be supported in individual cases (e.g., *Mimulus*; Brandvain et al., 2014)—testing the generality of these effects with a larger set of comparisons remains a goal in the future. Within the limitations of the data available here, however, mating system difference alone is an equivocal predictor of overall patterns of recent genomic introgression in this group.

Interestingly, we also did not detect an association between the magnitude of evolutionary divergence (genetic distance), and either the occurrence or the amount of inferred introgression. Species are expected to accumulate reproductive isolation with increasing evolutionary divergence (Coyne and Orr, 1997), and this pattern has been observed among wild tomatoes for loci involved in postzygotic reproductive isolation (hybrid pollen and seed sterility; Moyle and Nakazato, 2010), suggesting that introgression should become attenuated with increasing evolutionary age between species. Nonetheless, the total number of loci estimated to contribute to postzygotic isolation in this group is relatively modest, even among the oldest species pairs (Moyle and Nakazato, 2010, 2008). In addition, mean sequence divergence between all lineages analyzed here is low—0.1-4.0%—consistent with the recent, rapid origin of species in this clade (Pease et al., 2016). Using data from a very broad range of taxa, a recent meta-analysis inferred that divergence of just a few percent results in barriers that can effectively suppress gene flow (Roux et al., 2016) indicating that genetic divergence can be a strong determinant of introgression, at least beyond some threshold at which isolating barriers are sufficiently strong. The data presented here suggest that wild tomato species have not yet exceeded this threshold. Instead, our inference is that it is largely the opportunity for gene exchange, via geographical proximity, that currently determines the possibility of post-speciation introgression in this group.

Finally, as this inference suggests, the conditions that generally favor or prevent gene flow between species will likely vary depending on the biological features of different systems. Although there are yet too few comprehensive tests to determine how much they might vary, some previous analyses have assessed factors influencing general introgression patterns with different approaches. For example, Winger (2017) evaluated the relationship between introgression and plumage differentiation for 16 lineages of Andean cloud forest birds within a geographically and ecologically structured study. He found evidence for introgression across a geographic barrier between lineage pairs with uniform plumage patterns, but not between pairs with divergent plumage. This suggests that different patterns of sexual selection might determine whether and when introgression is expected, although alternative explanations, including more time since divergence between plumage-differentiated pairs, could not be excluded in this case. Clearly, additional systematic tests of the ecological, reproductive, and historical factors most strongly predictive of post-speciation gene flow will be helpful in evaluating how these might or might not differ between major groups of organisms.

### Evaluating introgression as an important evolutionary force

Our analyses join a growing consensus of studies that suggest gene flow among distinct lineages might be common, especially those that have rapidly diverged, have incomplete isolating barriers, and that maintain some regions of geographical overlap (Brawand et al., 2014; Jónsson et al., 2014; Lamichhaney et al., 2015; The Heliconius Genome Consortium et al., 2012). What do these observations indicate about whether introgression contributes substantially to shaping the evolution of these lineages? The expectations here are complex. When gene flow is frequent the opportunity for introgressed alleles to contribute to evolutionary responses in the recipient lineage clearly increases, but gene flow could also have deleterious consequences by introducing alleles that are maladaptive in this new background. Instead, large amounts of gene flow between species might indicate that introgressing loci have little or no detectable fitness effects: that is—that they are adaptively inconsequential. Conversely, even very restricted gene flow might still be consistent with the possibility of ‘adaptive’ introgression—the movement of alleles between species that increase fitness in their new recipient lineage (Suarez-Gonzalez et al., 2018). Several of the best cases of apparently adaptive introgression involve the movement of relatively small chromosomal segments (e.g. mimicry loci in *Heliconius*; high altitude adaptation in ancestral human populations; Suarez-Gonzalez et al., 2018; The Heliconius Genome Consortium et al., 2012). iven a genetic background of potentially deleterious interactions (that cause reduced hybrid fitness) a fairly limited exchange of alleles might be expected, especially if the associated variants have very large but local fitness consequences, as is observed for locally adaptive mimicry alleles in *Heliconius* for example. While not a primary goal of our analyses here, we did not observe any obvious analogous cases at mating-system loci in our genomes (see Supplementary Text). A prior analysis using transcriptome data did detect limited regions of introgression among some *Solanum* species pairs, with indirect evidence that these might have adaptive functions associated with local ecological conditions 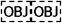 definitely demonstrating adaptive function requires knowing the fitness effects of alternative alleles under ecologically realistic conditions, these remain hypotheses. For now, the observation that a low level of gene exchange frequently occurs between lineages suggests that introgression could be a significant source of adaptive genetic variation, certainly in comparison to lineages where there is no evidence of gene flow.

Overall, while genetic exchange in any particular instance is likely to be influenced by both ecological and genomic contexts, only via systematic tests of introgression patterns across multiple cases can we start to disentangle the major determinants of post-speciation gene flow. One of our goals here was to demonstrate that structured *a priori* tests, across multiple species pairs in a diverse clade, provides one method for assessing the relative influence of several general factors on the frequency and amount of post-speciation introgression. By going beyond individual cases to generate general and quantitative evaluations, these tests should provide insight both into the main determinants of post-speciation introgression across a diversity of organisms and contexts, and the relative importance of introgression as an engine of evolution.

## Supporting information

Supplemental Methods and Results

## ACKNOWLEDGEMENTS

The authors thank members of the Moyle Lab and the Hahn Lab (Indiana University) for advice on genomic analyses, in addition to M. Behringer, J. Pease, and M. Hahn. The authors would also like to thank # anonymous reviewers for comments that greatly improved the manuscript. This research was funded by National Science Foundation grant MCB-1127059.

## AUTHOR CONTRIBUTIONS

J.A.P.H. and L.C.M. designed the experiments; J.A.P.H. conducted the bioinformatic analyses; J.A.P.H. and L.C.M. wrote the paper.

## DATA ACCESSIBILITY

Data used in the analyses are available from the NCBI SRA (Wild accessions of *Solanum* section *Lycopersicum*: SRP045767; *Solanum tuberosum*: SRP059592) or the European Nucleotide Archive (PRJEB5235). Output files generated by mvftools available via data dryad (XXXX).

