## Supplemental Methods and Results for "Spatial proximity determines post-speciation introgression in Solanum"

*Mapping raw reads to reference genome*

To combine the data from three different sequencing genome sequencing projects, we trimmed and re-mapped raw reads back to the reference genome of tomato, *S. lycopersicum* version 2.50 (The Tomato Genome Consortium, 2012). Trimmomatic was first used for removing adaptors and quality trimming, using a 4-base sliding window, and cutting when the average quality per base dropped below 20 and/or when reads were less than 36 bp long (Bolger et al., 2014). All retained reads were then mapped back to the reference genome using bwa mem (Li, 2013), sorted and indexed with samtools (Li et al., 2009), *mpileup* was used to generate a BCF file for each of the 12 chromosomes (to facilitate downstream analyses) for all accessions and then converted to VCF via bcftools. We then filtered the data using VCFtools such that the minimum base quality was required to be > 30 and read depth >10 generating a filtered VCF file for each chromosome, in which all base calls that did not pass quality filtering were then masked using the mvf_filter script. The combined and filtered VCF files were converted to mvf format using MVFtools (Pease and Rosenzweig, 2018) prior to calculating introgression statistics. For each 100kb window, a z-score was calculated to determine if the D value in this specific window was different from zero; individual 100kb windows with D significantly >0 are shown as colored points while non-significant windows are represented as black (Supplemental Figure 3) or grey (Figure 3 and 4) circles.

*Defining and determining geographical proximity, mating system difference, and evolutionary distance*

To test if spatial proximity increases the incidence of hybridization, we defined geographically closer lineages (P2 and P3) and geographically distant lineages (P1 and P3) based on known geographical ranges for our species and physical collection locations of the sequenced accessions. For these analyses, we only included species pairs that are known to be geographically proximate within their native ranges. To identify these species pairs, we used the collection location data (at tgrc.ucdavis.edu) to determine the geographical location of all accession records for each species, and then identified the subset for which the geographic distance between at least one accession of each species pairing was less than 5 km. Five km was chosen as a reasonable maximum cutoff for defining geographical proximity because a number of ecological studies, including in *Solanum*, suggest it is within potential dispersal distances for bee pollinators (Rick, 1950). Within this set of species pairs, we then identified appropriate accessions from our 32 sequenced accessions to use as P1, P2, and P3 in our 4-taxon tests.

For P2 and P3, we identified the geographically closest accessions for two species, based on inferred pairwise geographic distances among all sequenced accessions. In addition, we required that the P2 accession must be located within the known range of geographical overlap with the P3 species. To define a geographically distant (P1) accession to contrast with P2 from the same species, we chose the accession with the largest distance from the focal P3 accession, with a minimum geographical distance of at least 100km, from among our 32 sequenced accessions. Note that this P1 lineage could occur within the geographical range of overlap with the second species, but the specific P1 and P3 accessions must be greater than 100 km from each other.

From these criteria, 14 different comparisons (4-taxon tests) could be made (Table 1 and Supplemental Table 2). Because the actual geographic distances among these available accessions varies widely, this analysis is an imperfect reflection of close spatial proximity (sympatry); however, the structure of the tests means we are still systematically polarizing our comparisons to assess the effect of more (closer) versus less (distant) geographic proximity. We also directly considered quantitative differences in geographical proximity between P1 and P2 with accessions from the P3 species in our analyses.

For mating system tests, there are three species within *Solanum* in which both SI and SC populations are described and for which we had whole genome sequence data (*S. arcanum*, *S. habrochaites*, or *S. peruvianum*), allowing us to test if introgression with a second species is more likely given mating system similarities in these three cases (Table 1). For each 4-taxon test, the P1 and P2 position are occupied by self-incompatible and self-compatible accessions, respectively, of one species, while the P3 position is an accession from a self-compatible lineage (Figure 1). Given constraints on the structure of the test and available genome-sequence data, we used the same (*S. pimpinefollium* LA0373 or LA0400) accession in the P3 position in each of our three tests (see Results).

To evaluate whether D statistics change systematically over increasing evolutionary divergence, we estimated overall genetic divergence between species in each trio within each 4-taxon test by averaging the two estimates of pairwise genetic distance between P1 and P3 and between P2 and P3. Genetic distance was calculated as the total number of sites that differed divided by total number of sites for each pairwise comparison. Our estimates of genetic distance ranged from 0.3% to 4.0% (see Results, Supplementary Table 4) which is within the range expected for pairwise genome-wide divergence in this group (Aflitos et al., 2014; Lin et al., 2014; Pease et al., 2016). We then assessed the relationship between our estimate of mean genetic distance and the mean genome-wide estimate of D, across our assessed trios.

*Examining the association between introgression patterns and loci known to be associated with interspecific pollen-pistil barriers*

One expectation of mating system comparisons is that introgression patterns along the genome could be influenced by the genomic location of loci known to affect mating-system-related crossing compatibility. In the Solanaceae, several chromosomal regions have previously been associated both with the expression of gametophytic self-incompatibility and with pollen-pistil barriers between species (in particular, pistil-side barriers in which SI species reject pollen from SC species or ‘unilateral incompatibility’ (UI)) (Bernacchi and Tanksley, 1997; Li and Chetelat, 2014). In wild Solanum, these include loci on chromosome 1 (the S-locus itself), 12 (likely HT protein; Tovar-Méndez et al., 2017) and 3 (an unidentified gene(s) that magnifies the expression of pistil-side interspecific pollen rejection; Hamlin et al., 2017). Therefore, we further examined whether patterns of introgression along chromosomes 1, 3, and 12 in our mating system analysis were associated with these regions. To do so, we generated a list of genes located in chromosomal blocks that showed highly significant D-statistics in our analyses (Supplementary Figure 4). First, we identified the genomic coordinates along each chromosome at which there was a shift to a block in which there was overrepresentation of the alternative (minority) topology (i.e. higher counts in ABBA or BABA vs. BBAA); we considered a ‘block’ to be a location in which there were at least ten consecutive 100 kb windows which demonstrated an overrepresentation of either ABBA or BABA sites. Using these genomic coordinates, *Bedtools* (Quinlan and Hall, 2010), and the *ITAG2.4* gene models, we extracted all known genes located within the introgressed blocks, including genes which were up to 50 kb upstream and downstream from the endpoints of each estimated introgressed block (Supplementary Table 9). We found no evidence that chromosomal blocks with highly significant D statistics were associated with loci previously implicated in intra- (SI) and inter-specific (UI) pistil-side mating barriers (Supplemental Table 9a - i).

Supplemental Table 1. Species, accession, associated ID for the three different sequencing projects and the species used in this analysis.

| Species | Accession | Unique ID* | Sequencing Project |
| --- | --- | --- | --- |
| *S. corneliomulleri* | LA0118 | 25 | Tomato.150 |
| *S. pimpinellifolium* | LA0373 | SRR1572235 | Tomato.360 |
| *S. pimpinellifolium* | LA0400 | SRR1572257 | Tomato.360 |
| *S. habrochaites* | LA0407 | 71 | Tomato.150 |
| *S. pimpinellifolium* | LA0417 | SRR1572243 | Tomato.360 |
| *S. pimpinellifolium* | LA0442 | SRR1572238 | Tomato.360 |
| *S. cheesmaniae* | LA0483 | 53 | Tomato.150 |
| *S. pennelli* | LA0716 | 74 | Tomato.150 |
| *S. cheesmaniae* | LA0746 | SRR1572688 | Tomato.360 |
| *S. galapagense* | LA1044 | 104 | Tomato.150 |
| *S. pimpinellifolium* | LA1245 | SRR1572248 | Tomato.360 |
| *S. pimpinellifolium* | LA1246 | SRR1572234 | Tomato.360 |
| *S. pimpinellifolium* | LA1269 | SRR1572240 | Tomato.360 |
| *S. peruvianum* | LA1278 | 49 | Tomato.150 |
| *S. pimpinellifolium* | LA1341 | SRR1572270 | Tomato.360 |
| *S. huaylense* | LA1365 | 63 | Tomato.150 |
| *S. pimpinellifolium* | LA1375 | SRR1572239 | Tomato.360 |
| *S. pimpinellifolium* | LA1521 | SRR1572241 | Tomato.360 |
| *S. pimpinellifolium* | LA1582 | SRR1572246 | Tomato.360 |
| *S. pimpinellifolium* | LA1595 | SRR1572288 | Tomato.360 |
| *S. pimpinellifolium* | LA1617 | SRR1572250 | Tomato.360 |
| *S. habrochaites* | LA1718 | 69 | Tomato.150 |
| *S. habrochaites* | LA1777 | 70 | Tomato.150 |
| *S. pimpinellifolium* | LA1933 | SRR1572273 | Tomato.360 |
| *S. peruvianum* | LA1954 | 60 | Tomato.150 |
| *S. chilense* | LA1969 | SRR1572696 | Tomato.360 |
| *S. huaylense* | LA1983 | 62 | Tomato.150 |
| *S. neorickii* | LA2133 | 56 | Tomato.150 |
| *S. pimpinellifolium* | LA2147 | SRR1572274 | Tomato.360 |
| *S. arcanum* | LA2157 | 58 | Tomato.150 |
| *S. arcanum* | LA2172 | 59 | Tomato.150 |
| *S.peruvianum* | PI126935 | SRR1572694 | Tomato.360 |
| *S. tuberosum* | - | SRR2069932 | Potato |

*SRR unique IDs are from NCBI SRA website; numeric unique IDs are from

http://www.tomatogenome.net/accessions.html.

Supplemental Table 2. Pairwise geographic distances among P1, P2, and P3 accessions used in all analyzed trios (4-taxon tests). The trio name in the table represents the order of accessions in P1, P2, and P3 position. In all instances, we use the potato genome (*S. tuberosum*) as the outgroup.

| **Species in Trio** | **Accessions in Trio** | **Geo Dis**  **P1 and P2** | | | **Geo Dis**  **P1 and P3** | | **Geo Dis P2 and P3** |
| --- | --- | --- | --- | --- | --- | --- | --- |
| *Geographic trios* | | | | | | | |
| gal.gal.che | LA1044.LA0483.LA0746 | | 112 | 125 | | 72 | |
| arc.arc.pim | LA2172.LA2157.LA2147 | | 57 | 133 | | 79 | |
| pim.pim.neo | LA1375.LA1246.LA2133 | | 811 | 1077 | | 68 | |
| pim.pim.chi | LA1582.LA1933.LA1969 | | 1180 | 1645 | | 526 | |
| pim.pim.cor.1 | LA0400.LA1269.LA0118 | | 756 | 781 | | 50 | |
| pim.pim.cor.2 | LA1617.LA1521.LA0118 | | 1107 | 973 | | 141 | |
| pim.pim.per.1 | LA1595.LA1341.LA1278 | | 350 | 310 | | 234 | |
| pim.pim.per.2 | LA1617.LA1269.LA1278 | | 950 | 973 | | 25 | |
| pim.pim.hab | LA0417.LA0442.LA1777 | | 786 | 794 | | 65 | |
| pim.pim.pen | LA1245.LA1269.LA1272 | | 941 | 950 | | 13 | |
| arc.arc.hab | LA2172.LA2157.LA1718 | | 57 | 416 | | 359 | |
| hab.hab.neo | LA1777.LA1718.LA2133 | | 516 | 700 | | 205 | |
| hab.hab.cor | LA0407.LA1777.LA0118 | | 851 | 1091 | | 243 | |
| hua.hua.hab | LA1983.LA1365.LA1718 | | 173 | 101 | | 71 | |
| *Mating System Trios* | | | | | | | |
| arcSI.arcSC.pim.SC | LA2172.LA2157.LA0373 | | 57 | 442 | | 385 | |
| arcSI.arcSC.pim.SCv2 | LA2172.LA2157.LA0400 | | 57 | 143 | | 188 | |
| habSI.habSC.pimSC | LA1777.LA0407.LA0373 | | 851 | 75 | | 877 | |
| habSI.habSC.pimSCv2 | LA1777.LA0407.LA0400 | | 851 | 538 | | 341 | |
| perSI.perSC.pimSC | LA1278.PI128650.LA0373 | | 1053 | 234 | | 1286 | |
| perSI.perSC.pimSCv2 | LA1278.PI128650.LA0400 | | 1053 | 779 | | 1811 | |

Supplemental Table 3. Number of windows examined for each trio for both the entire dataset and the reduced dataset (i.e. only windows for which number of informative sites were greater than 20).

| Species in Trio | Accession in Trio | Full dataset | Reduced dataset |
| --- | --- | --- | --- |
| *Geographic Comparisons* |  |  |  |
| gal.gal.che | LA1044.LA0483.LA0746 | 8027 | 150 |
| arc.arc.pim | LA2172.LA2157.LA2147 | 8027 | 5333 |
| pim.pim.neo | LA1375.LA1246.LA2133 | 8027 | 5224 |
| pim.pim.chi | LA1582.LA1933.LA1969 | 8027 | 3141 |
| pim.pim.cor1 | LA0400.LA1269.LA0118 | 8027 | 4381 |
| pim.pim.cor2 | LA1617.LA1521.LA0118 | 8027 | 4205 |
| pim.pim.per1 | LA1595.LA1341.LA1278 | 8027 | 5710 |
| pim.pim.per2 | LA1617.LA1269.LA1278 | 8027 | 4263 |
| pim.pim.hab | LA0417.LA0442.LA1777 | 8027 | 5416 |
| pim.pim.pen | LA1245.LA1269.LA1272 | 8027 | 3358 |
| arc.arc.hab | LA2172.LA2157.LA1718 | 8027 | 5129 |
| hab.hab.neo | LA1777.LA1718.LA2133 | 8027 | 5223 |
| hab.hab.cor | LA0407.LA1777.LA0118 | 8027 | 5451 |
| hua.hua.hab | LA1983.LA1365.LA1718 | 8027 | 4876 |
| *Mating System Comparisons* |  |  |  |
| arcSI.arcSC.pim.SC | LA2172.LA2157.LA0373 | 8027 | 5234 |
| habSI.habSC.pimSC | LA1777.LA0407.LA0373 | 8027 | 5800 |
| perSI.perSC.pimSC | LA1278.PI128650.LA0373 | 8027 | 4182 |

Supplemental Table 4. Introgression statistics for each trio, using the dataset that includes all windows regardless of the number of informative sites. All trios are significantly different than zero after Bonferroni correction. For each trio, the order in which each species is listed corresponds to (P1, P2, P3); accessions are the same as Table 1 and are listed in the same order.

| Group | Accessions in Trio | MeanD | StnDev | SE | C95_lwr | C95_upr | chi square | p.value |
| --- | --- | --- | --- | --- | --- | --- | --- | --- |
| *Geographic Trios* | | | | | | | |  |
| gal.gal.che | LA1044.LA0483.LA0746 | -0.027 | 0.488 | 0.005 | -0.038 | -0.017 | 93.96 | 2.81E-24 |
| arc.arc.pim | LA2172.LA2157.LA2147 | 0.024 | 0.535 | 0.005 | 0.013 | 0.034 | 1819.01 | 5.64E-230 |
| pim.pim.neo | LA1375.LA1246.LA2133 | -0.081 | 0.522 | 0.005 | -0.091 | -0.070 | 146.57 | 5.20E-23 |
| pim.pim.chi | LA1582.LA1933.LA1969 | 0.008 | 0.372 | 0.005 | -0.002 | 0.019 | 20.94 | 4.20e-18 |
| pim.pim.cor1 | LA0400.LA1269.LA0118 | 0.015 | 0.404 | 0.005 | 0.005 | 0.026 | 21.23 | 2.50E-04 |
| pim.pim.cor2 | LA1617.LA1521.LA0118 | 0.090 | 0.535 | 0.005 | 0.079 | 0.100 | 244.38 | 3.44E-28 |
| pim.pim.per1 | LA1595.LA1341.LA1278 | 0.007 | 0.440 | 0.005 | -0.004 | 0.017 | 26.12 | 5.10E-05 |
| pim.pim.per2 | LA1617.LA1269.LA1278 | 0.069 | 0.506 | 0.005 | 0.059 | 0.080 | 154.19 | 3.22E-19 |
| pim.pim.hab | LA0417.LA0442.LA1777 | 0.031 | 0.491 | 0.005 | 0.021 | 0.042 | 114.62 | 1.50E-21 |
| pim.pim.pen | LA1245.LA1269.LA1272 | 0.060 | 0.414 | 0.005 | 0.049 | 0.070 | 179.11 | 1.90E-16 |
| arc.arc.hab | LA2172.LA2157.LA1718 | -0.019 | 0.455 | 0.005 | -0.030 | -0.009 | 33.929 | 5.11E-98 |
| hab.hab.neo | LA1777.LA1718.LA2133 | 0.036 | 0.487 | 0.005 | 0.025 | 0.046 | 55.48 | 7.34E-26 |
| hab.hab.cor | LA0407.LA1777.LA0118 | 0.043 | 0.473 | 0.005 | 0.032 | 0.053 | 168.95 | 2.2e-16 |
| hua.hua.hab | LA1983.LA1365.LA1718 | 0.037 | 0.468 | 0.005 | 0.026 | 0.047 | 393.008 | 8.54E-12 |
| *Mating system trios* | | | | | | | |  |
| arcSI.arcSC.pimSC | LA2172.LA2157.LA0373 | 0.0278 | 0.5325 | 0.0057 | 0.0165 | 0.0390 | 1856.19 | 0 |
| arcSI.arcSC.pimSCv2 | LA2172.LA2157.LA0400 | 0.0235 | 0.542 | 0.0058 | 0.0120 | 0.0349 | 835.46 | 3.15E-183 |
| habSI.habSC.pimSC | LA1777.LA0407.LA0373 | -0.0293 | 0.4878 | 0.0057 | -0.0406 | -0.0181 | 243.63 | 1.92E-54 |
| habSI.habSC.pimSCv2 | LA1777.LA0407.LA0400 | -0.0250 | 0.5142 | 0.0058 | -0.0364 | -0.0131 | 72.88 | 4.08E-17 |
| perSI.perSC.pimSC | LA1278.PI128650.LA0373 | 0.0932 | 0.5174 | 0.0057 | 0.0820 | 0.1045 | 5578.46 | 2.2e-16 |
| perSI.perSC.pimSCv2 | LA1278.PI128650.LA0400 | 0.0913 | 0.5073 | 0.0058 | 0.0798 | 0.1027 | 3275.12 | 2.2e-16 |

StnDev = standard deviation; SE = standard error of mean; C95_lwr and C95_upr = Lower and upper 95% confidence intervals, respectively, Chi-square = Chi-squared goodness-of-fit test value; p.value* = p value adjusted after Bonferroni correction. When the entire dataset is included, our empirical estimates for genome-wide D-statistics are similar to our estimates from only windows with > 20 sites, although the estimated magnitude of mean D decreased (i.e. was closer to zero) for some trios.

Supplemental Table 5a. Geographic distance between either P1 or P2 and their closet accession from the species used in the P3 position. The relative difference is calculated as: (P1, P3) – (P2, P3).

| Species in Trio | Accessions in Trio | P1 or P2 Accession | | Closest P3 Accession | | Geographic  distance (km) | | Relative Geographic between P1/P3 and P2/P3 |
| --- | --- | --- | --- | --- | --- | --- | --- | --- |
| *Geographic trios* |  |  | |  | |  | |  |
| gal.gal.che | LA1044.LA0483.LA0746 | LA1044 | | LA0425 | | 28 | | 28 |
| gal.gal.che | LA1044.LA0483.LA0746 | LA0483 | | LA0521 | | 0 | |  |
| arc.arc.pim | LA2172.LA2157.LA2147 | LA2172 | | LA0398 | | 48 | | 20 |
| arc.arc.pim | LA2172.LA2157.LA2147 | LA2157 | | LA2181 | | 28 | |  |
| pim.pim.neo | LA1375.LA1246.LA2133 | LA1375 | | LA2613 | | 202 | | 194 |
| pim.pim.neo | LA1375.LA1246.LA2133 | LA1246 | | LA2113 | | 8 | |  |
| pim.pim.chi | LA1582.LA1933.LA1969 | LA1582 | | LA1917 | | 1041 | | 1038 |
| pim.pim.chi | LA1582.LA1933.LA1969 | LA1933 | | LA1934 | | 3 | |  |
| pim.pim.cor.1 | LA0400.LA1269.LA0118 | LA0400 | | LA3664 | | 35 | | 32 |
| pim.pim.cor.1 | LA0400.LA1269.LA0118 | LA1269 | | LA1271 | | 3 | |  |
| pim.pim.cor.2 | LA1617.LA1521.LA0118 | LA1617 | | LA1379 | | 895 | | 894 |
| pim.pim.cor.2 | LA1617.LA1521.LA0118 | LA1521 | | LA1609 | | 1 | |  |
| pim.pim.per.1 | LA1595.LA1341.LA1278 | LA1595 | | LA0372 | | 79 | | 78 |
| pim.pim.per.1 | LA1595.LA1341.LA1278 | LA1341 | | LA0370 | | 1 | |  |
| pim.pim.per.2 | LA1617.LA1269.LA1278 | LA1617 | | LA2849 | | 303 | | 303 |
| pim.pim.per.2 | LA1617.LA1269.LA1278 | LA1269 | | LA1270 | | 0 | |  |
| pim.pim.hab | LA0417.LA0442.LA1777 | LA0417 | | LA0407 | | 61 | | 20 |
| pim.pim.hab | LA0417.LA0442.LA1777 | LA0442 | | LA1361 | | 41 | |  |
| pim.pim.pen | LA1245.LA1269.LA1272 | LA1245 | | LA2654 | | 237 | | 224 |
| pim.pim.pen | LA1245.LA1269.LA1272 | LA1269 | | LA0751 | | 13 | |  |
| arc.arc.hab | LA2157.LA2172.LA1777 | LA2157 | | LA2156 | | 3 | | 47 |
| arc.arc.hab | LA2157.LA2172.LA1777 | LA2172 | | LA2159 | | 50 | |  |
| hab.hab.neo | LA1777.LA1718.LA2133 | LA1777 | | LA0247 | | 122 | | 121 |
| hab.hab.neo | LA1777.LA1718.LA2133 | LA1718 | | LA1716 | | 1 | |  |
| hab.hab.cor | LA0407.LA1777.LA0118 | LA0407 | | LA1379 | | 1013 | | 848 |
| hab.hab.cor | LA0407.LA1777.LA0118 | LA1777 | | LA1379 | | 165 | |  |
| hua.hua.hab | LA1983.LA1365.LA1777 | LA1983 | | LA1393 | | 44 | | 43 |
| hua.hua.hab | LA1983.LA1365.LA1777 | LA1365 | | LA1366 | | 1 | |  |
| *Mating system trios* |  | |  | |  | |  | |
| arcSI.arcSC.pim.SC | LA2172.LA2157.LA0373 | LA2172 | | LA2181 | | 28 | | 20 |
| arcSI.arcSC.pim.SC | LA2172.LA2157. LA0373 | LA2157 | | LA0398 | | 48 | |  |
| habSI.habSC.pimSC | LA1777.LA0407. LA0373 | LA1777 | | LA2575 | | 19 | | 17 |
| habSI.habSC.pimSC | LA1777.LA0407. LA0373 | LA0407 | | LA1399 | | 2 | |  |
| perSI.perSC.pimSC | LA1278. PI128650*. LA0373 | LA1278 | | LA1277 | | 0 | | 83 |
| perSI.perSC.pimSC | LA1278. PI128650*. LA0373 | PI128650* | | LA1670 | | 83 | |  |

*LA1537 was used as a proxy for PI128650; this GPS coordinate is an estimate based on LA1537 which it was collected in the Azapa Valley and is about 3 km from Arica, Chile (https://cgngenis.wur.nl/AccessionDetails.aspx?acnumber=CGN15876)

Supplemental Table 6. The proportion of the genome estimated to have differentially experienced introgression in each trio, calculated as (ABBA-BABA)/ (ABBA + BABA + BBAA). Trios are listed in the same order as Table 1.

| Species in Trio | Sum  BBAA | Sum  ABBA | Sum  BABA | Est. proportion of genome differentially introgressed |
| --- | --- | --- | --- | --- |
| *Geographic trios* |  |  |  |  |
| gal.gal.che | 5693 | 120 | 146 | -0.0044 |
| arc.arc.pim | 485851 | 113346 | 102512 | 0.0154 |
| pim.pim.neo | 553452 | 7444 | 9094 | -0.0029 |
| pim.pim.chi | 154067 | 1019 | 971 | 0.0003 |
| pim.pim.cor1 | 328264 | 1405 | 1237 | 0.0005 |
| pim.pim.cor2 | 291137 | 3921 | 2773 | 0.0039 |
| pim.pim.per1 | 869492 | 3048 | 2697 | 0.0004 |
| pim.pim.per2 | 322514 | 3364 | 2547 | 0.0025 |
| pim.pim.hab | 768289 | 2992 | 2718 | 0.0004 |
| pim.pim.pen | 167869 | 1266 | 798 | 0.0028 |
| arc.arc.hab | 794821 | 70780 | 73774 | -0.0032 |
| hab.hab.neo | 926704 | 29465 | 26532 | 0.0030 |
| hab.hab.cor | 941988 | 38888 | 33712 | 0.0051 |
| hua.hua.hab | 383617 | 56852 | 50018 | 0.0139 |
| *Mating System trios* |  |  |  |  |
| arcSI.arcSC.pimSC | 445093 | 105201 | 93846 | 0.0176 |
| arcSI.arcSC.pimSCv2 | 240809 | 56651 | 51951 | 0.0135 |
| habSI.habSC.pimSC | 930630 | 37611 | 37964 | -0.0004 |
| habSI.habSC.pimSCv2 | 517957 | 20932 | 21011 | -0.0001 |
| perSI.perSC.pimSC | 164127 | 31101 | 15197 | 0.0756 |
| perSI.perSC.pimSCv2 | 82102 | 15451 | 7352 | 0.0772 |

Supplemental Table 8. Estimated genome-wide genetic distance (expressed as percent sequence difference) for each geographic and mating system trio. Trios are listed in the same order as Table 1.

| Species in Trio | Accessions in Trio | Gen.dis* P1 and P3 | | Gen.dis* P2 and P3 | | Averaged gen.dis* |
| --- | --- | --- | --- | --- | --- | --- |
| *Geographic trios* |  | | | | | |
| gal.gal.che | LA1044.LA0483.LA0746 | 0.24 | 0.44 | | 0.34 | |
| arc.arc.pim | LA2172.LA2157.LA2147 | 2.80 | 3.11 | | 2.95 | |
| pim.pim.neo | LA1375.LA1246.LA2133 | 2.96 | 2.93 | | 2.95 | |
| pim.pim.chi | LA1582.LA1933.LA1969 | 3.00 | 3.03 | | 3.02 | |
| pim.pim.cor1 | LA0400.LA1269.LA0118 | 3.58 | 3.62 | | 3.60 | |
| pim.pim.cor2 | LA1617.LA1521.LA0118 | 3.59 | 3.48 | | 3.53 | |
| pim.pim.per1 | LA1595.LA1341.LA1278 | 3.47 | 3.60 | | 3.54 | |
| pim.pim.per2 | LA1617.LA1269.LA1278 | 3.51 | 3.36 | | 3.44 | |
| pim.pim.hab | LA0417.LA0442.LA1777 | 3.87 | 3.80 | | 3.83 | |
| pim.pim.pen | LA1245.LA1269.LA1272 | 3.45 | 3.59 | | 3.52 | |
| arc.arc.hab | LA2172.LA2157.LA1718 | 3.67 | 3.87 | | 3.77 | |
| hab.hab.neo | LA1777.LA1718.LA2133 | 3.98 | 4.04 | | 4.01 | |
| hab.hab.cor | LA0407.LA1777.LA0118 | 3.72 | 3.76 | | 3.74 | |
| hua.hua.hab | LA1983.LA1365.LA1718 | 4.00 | 3.84 | | 3.92 | |
| *Mating System trios* |  | | | | | |
| arcSI.arcSC.pimSC | LA2172.LA2157.LA0373 | 2.92 | 2.85 | | 2.88 | |
| arcSI.arcSC.pimSCv2 | LA2172.LA2157.LA0400 | 2.911 | 2.88 | | 2.89 | |
| habSI.habSC.pimSC | LA1777.LA0407.LA0373 | 3.83 | 4.08 | | 3.95 | |
| habSI.habSC.pimSCv2 | LA1777.LA0407.LA0400 | 3.61 | 3.81 | | 3.71 | |
| perSI.perSC.pimSC | LA1278.PI128650.LA0373 | 3.65 | 2.82 | | 3.23 | |
| perSI.perSC.pimSCv2 | LA1278.PI128650.LA0400 | 3.477 | - | | 3.47 | |

*Gen.dis = Genome wide estimate of genetic distance as percentages.

Supplemental Figure Legends

Supplemental Figure 1. Relationship between genome-wide mean D and the relative difference in geographic distance of P1 and P2 from a heterospecific P3 population, for a) geographic trios, and b) mating system trios.

Supplemental Figure 2. Relationship between the amount of the genome estimated to have differentially experienced introgression (percent excess of one minority site pattern; see main text), and the average genome-wide genetic distance, for each analyzed trio.

Supplemental Figure 3. Chromosome-by-chromosome mean D, 95% CI, and distribution of D-statistic estimates from individual 100kb windows (black circles: window D not significantly different than zero; red circles: window D significantly different than zero), for each geographic trio. See Table 1 for abbreviations.

Supplemental Figure 4. The distribution of inferred tree topologies across the genome, for each trio in our mating system tests. Plots show the inferred topology for each 100kb window along the physical length of each chromosome. Genome-wide, the most probable topology is the species phylogeny (BBAA where (((P1, P2), P3), outgroup)). Alternative, ABBA (((P2, P3), P1), outgroup) or BABA (((P1, P3), P2), outgroup) topologies can be due to ILS or introgression. In our analysis, there is no clear association between genomic locations of known mating system loci, and the appearance of minority tree topologies, indicating that introgression is not unusually enriched or precluded specifically in these locations (see Supplemental Text).
