## Supplementary figures and images for "Spatial proximity determines post-speciation introgression in Solanum"

### Supplemental Figure 1

#### a. Geographic Trios

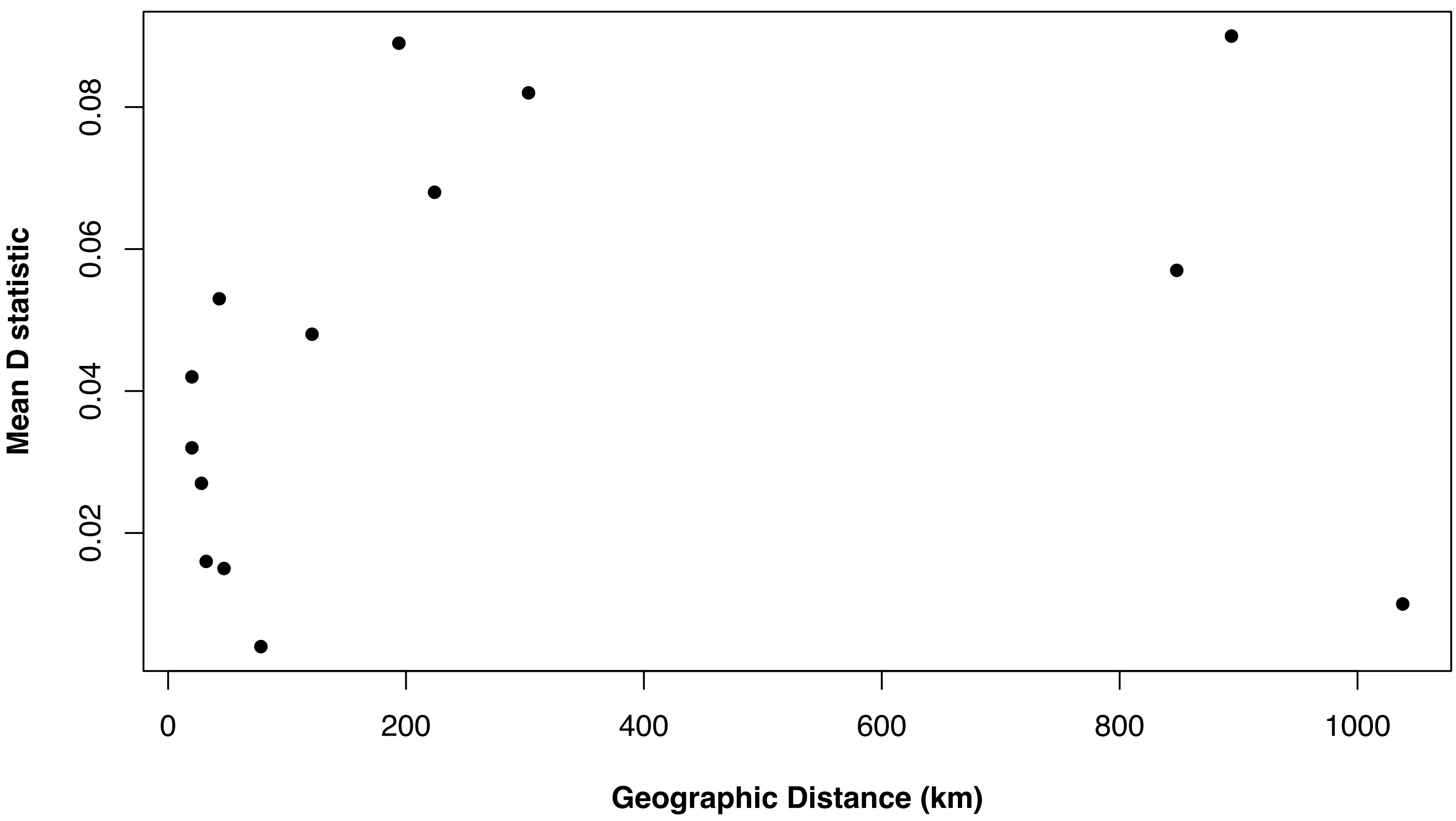


#### b. Mating System Trios

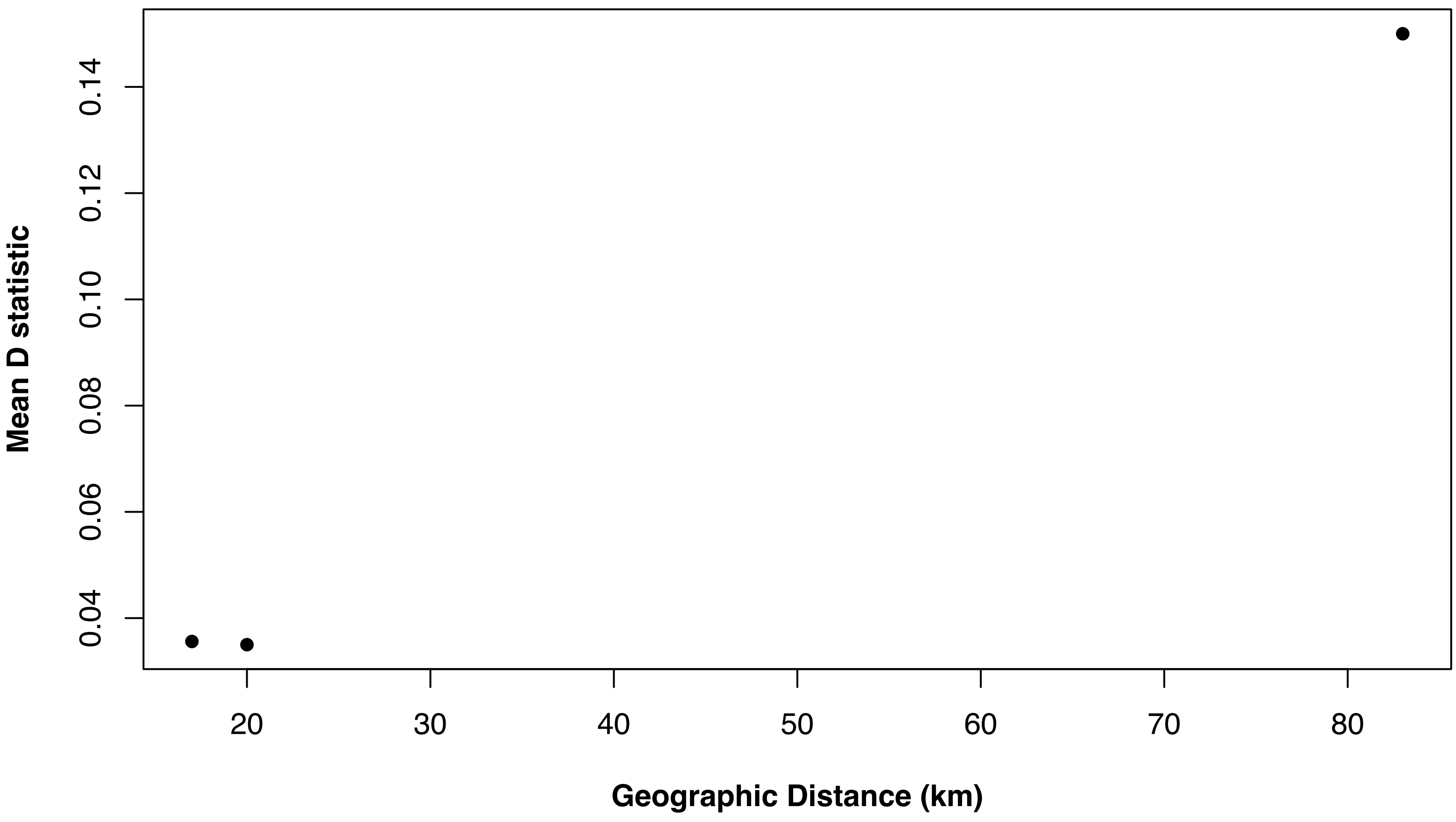

### Supplemental Figure 2

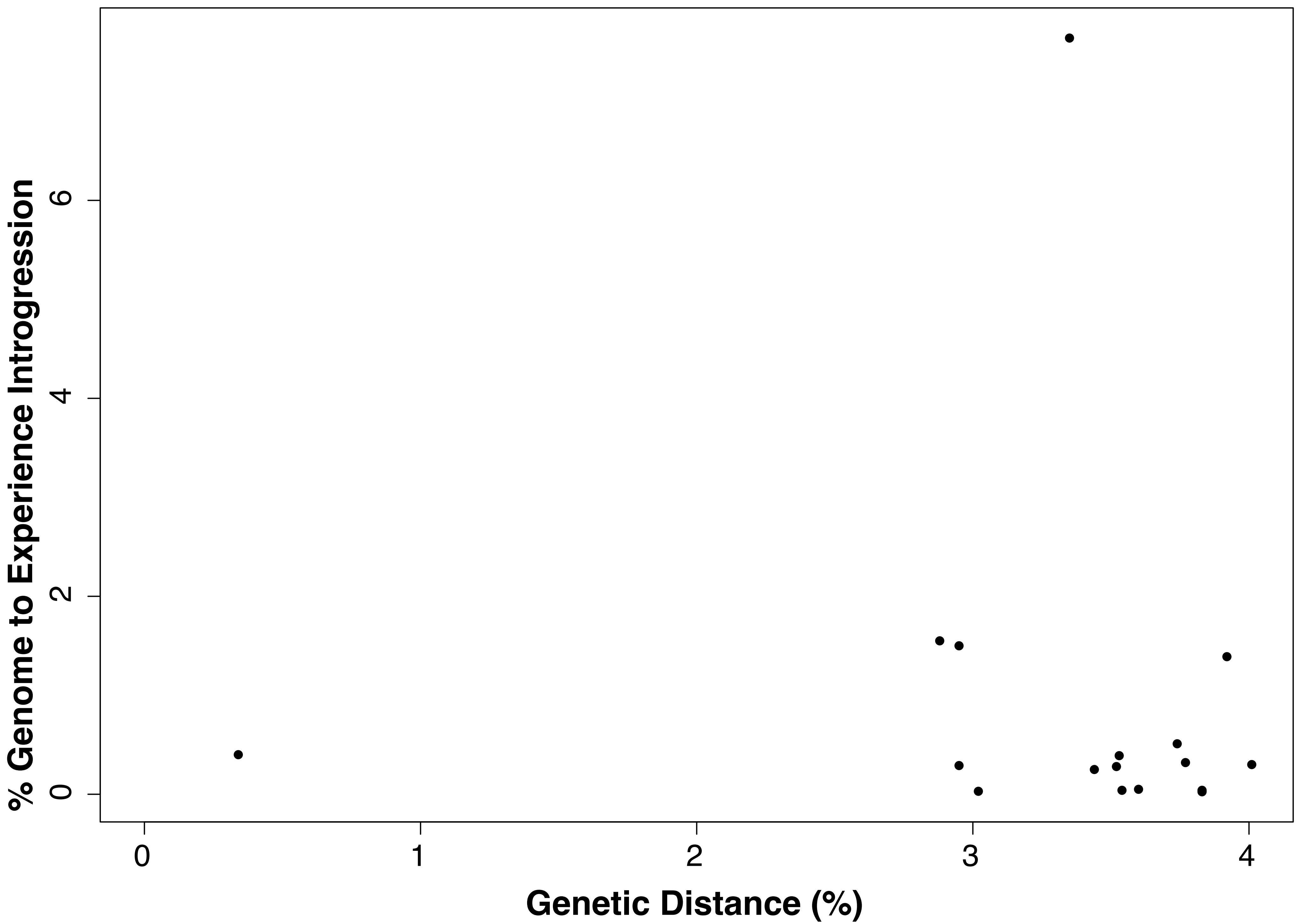

### Supplemental Figure 3

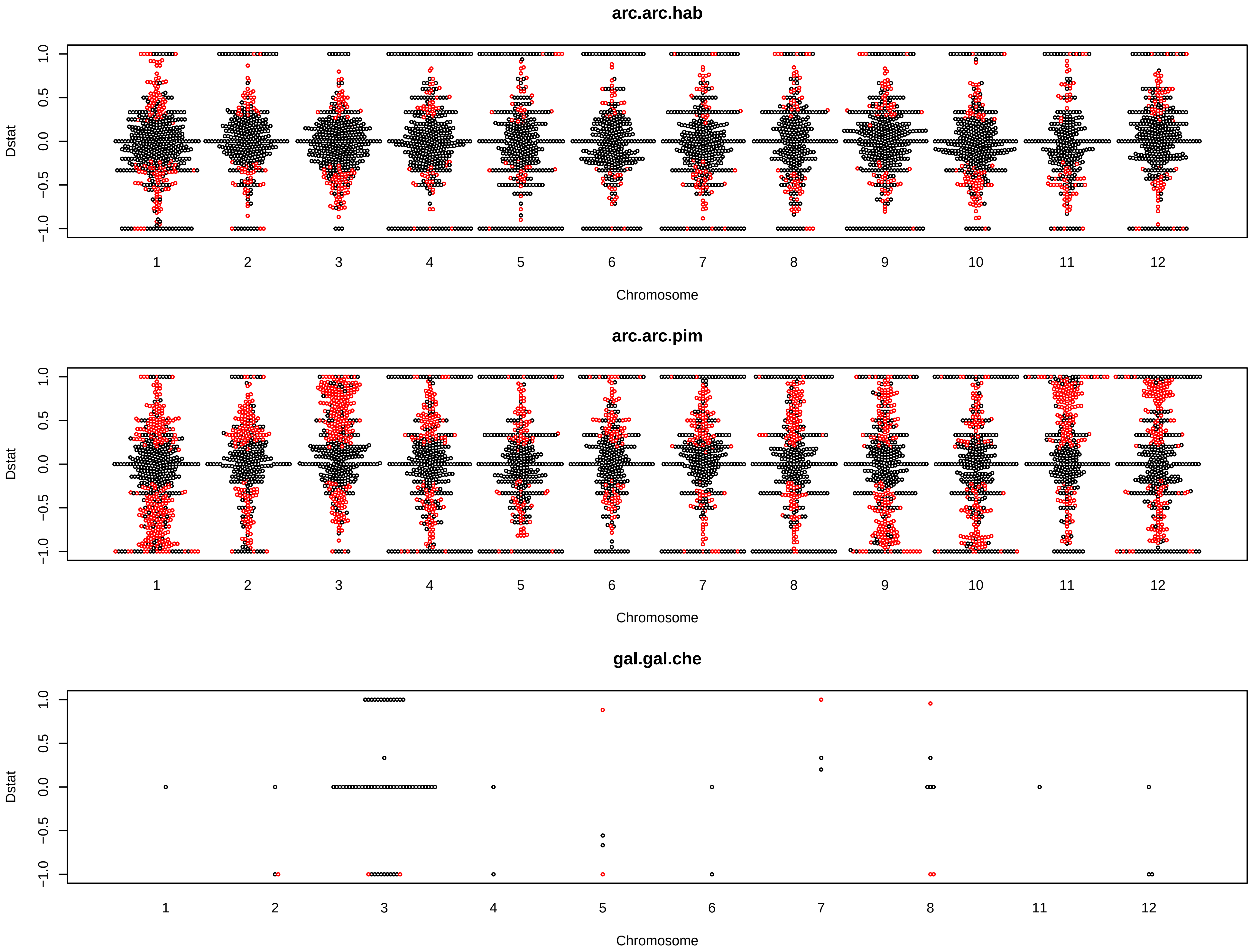

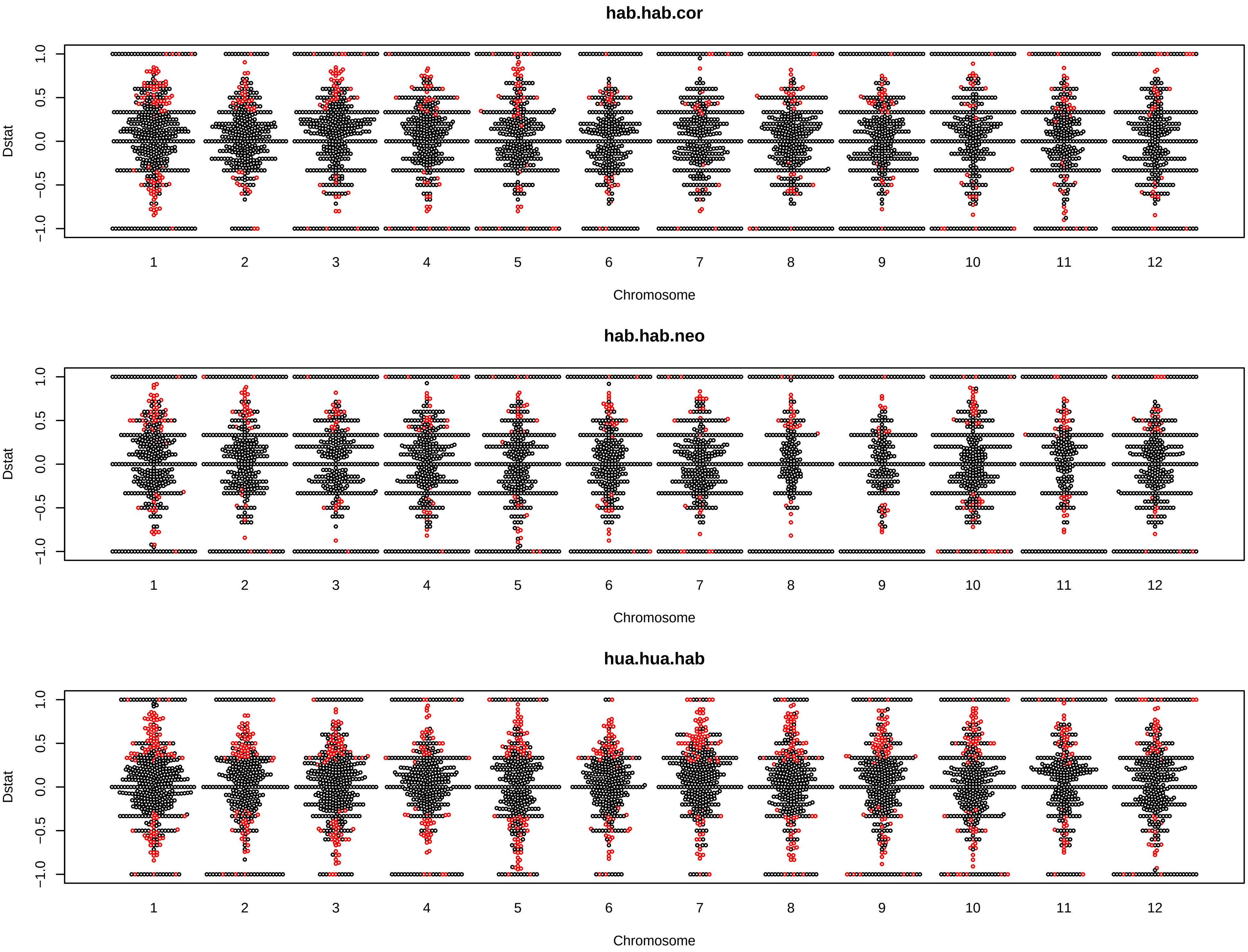

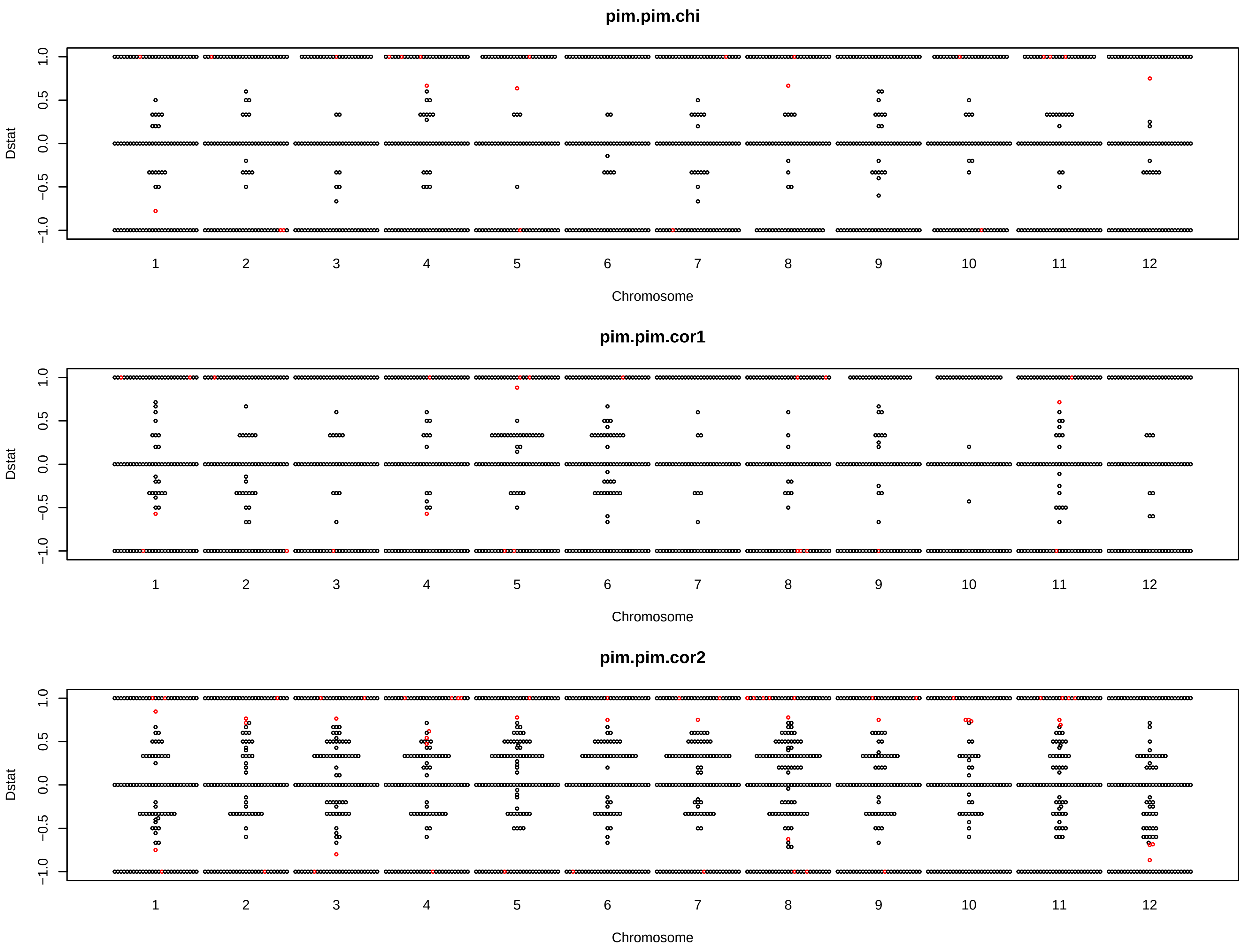

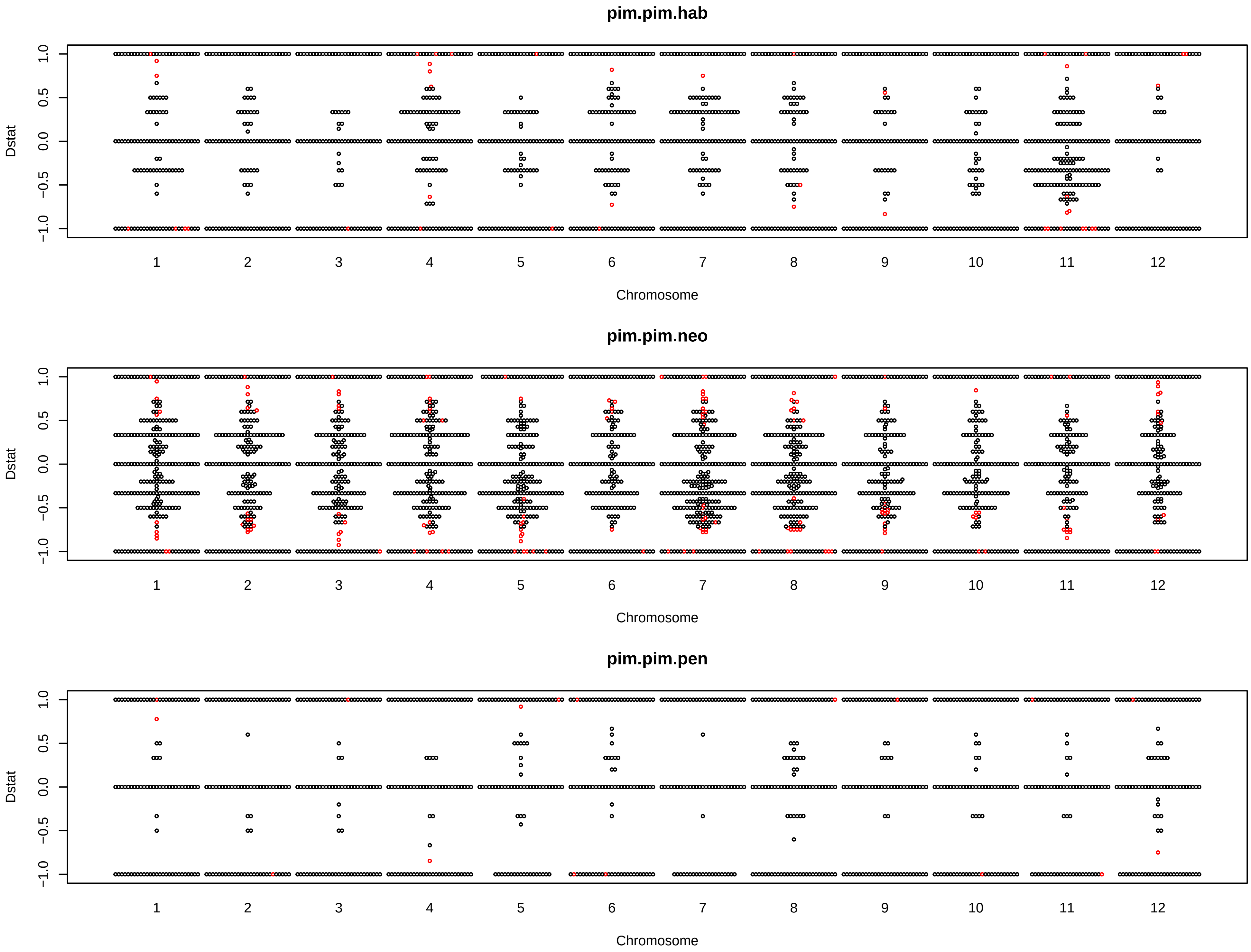

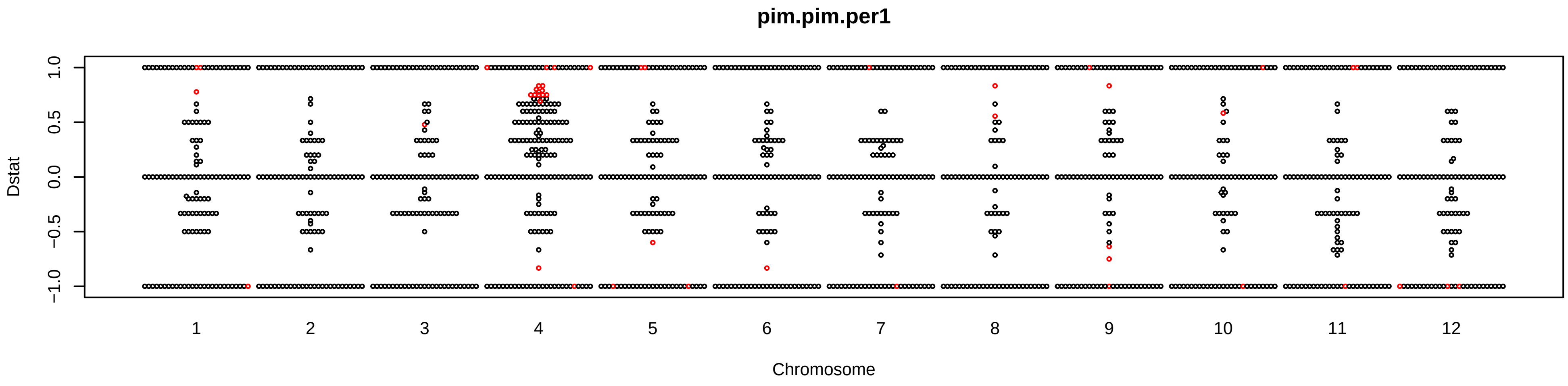

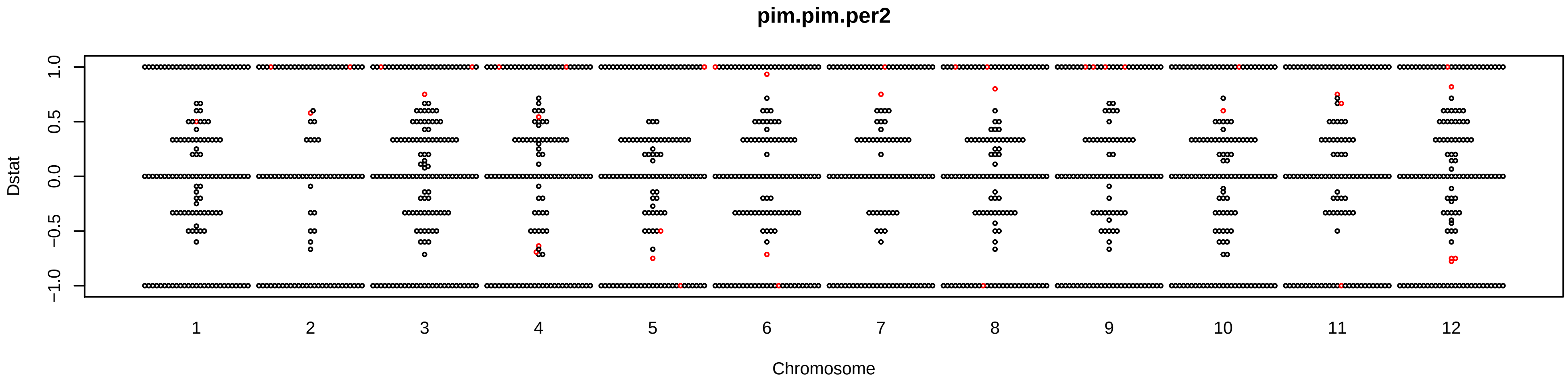

### Supplemental Figure 4

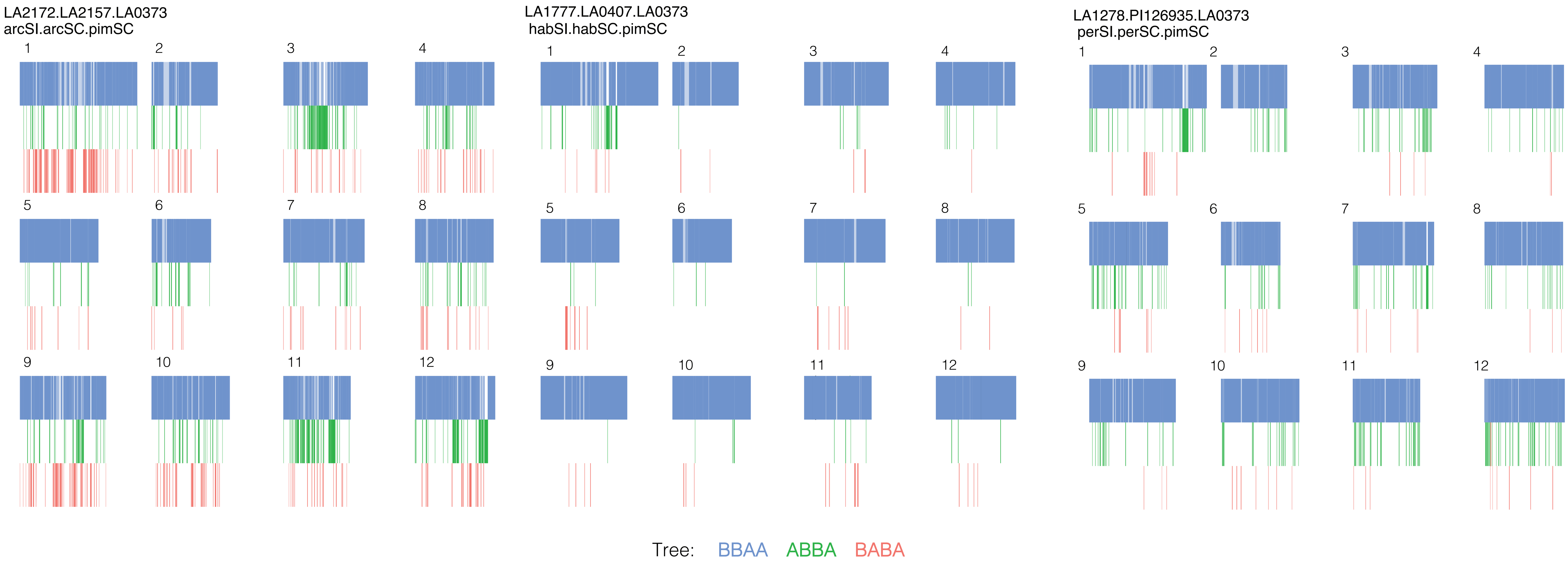
